# Distributed networks for auditory memory differentially contribute to recall precision

**DOI:** 10.1101/2021.01.18.427143

**Authors:** Sung-Joo Lim, Christiane Thiel, Bernhard Sehm, Lorenz Deserno, Jöran Lepsien, Jonas Obleser

**Author notes:** Correspondence: Sung-Joo Lim Department of Psychology, Binghamton University 4400 Vestal Parkway E Binghamton, NY 13902; Jonas Obleser Department of Psychology, University of Lübeck Maria-Goeppert-Str. 9a 23562 Lübeck, Germany.

## Abstract

Re-directing attention to objects in working memory can enhance their representational fidelity. However, how this attentional enhancement of memory representations is implemented across distinct, sensory and cognitive-control brain network is unspecified. The present fMRI experiment leverages psychophysical modelling and multivariate auditory- pattern decoding as behavioral and neural proxies of mnemonic fidelity. Listeners performed an auditory syllable pitch-discrimination task and received retro-active cues to selectively attend to a to-be-probed syllable in memory. Accompanied by increased neural activation in fronto-parietal and cingulo-opercular networks, valid retro-cues yielded faster and more perceptually sensitive responses in recalling acoustic detail of memorized syllables. Information about the cued auditory object was decodable from hemodynamic response patterns in superior temporal sulcus (STS), fronto-parietal, and sensorimotor regions. However, among these regions retaining auditory memory objects, neural fidelity in the left STS and its enhancement through attention-to-memory best predicted individuals’ gain in auditory memory recall precision. Our results demonstrate how functionally discrete brain regions differentially contribute to the attentional enhancement of memory representations.

## Introduction

Working memory is a short-term mental storage that maintains perceptual information even in the absence of sensory input. A hallmark of working memory is its limited capacity—that is, perceptual information held in memory is inherently noisy (Cowan, 2001; Bays and Husain, 2008; Luck and Vogel, 2013; Ma et al., 2014).

Selective attention plays a crucial role in overcoming the bottleneck of this limited capacity. Directing attention to the relevant information in memory can reduce inherent noise and enhance representations of the attended memory object—a process referred to as retrospective attention (Griffin and Nobre, 2003; Postle, 2006; Gazzaley and Nobre, 2012; Myers et al., 2017). Thus, even without sensory information, selective attention interacts with working memory and facilitates the maintenance of attended memory objects.

Research on the neural bases of working memory and selective attention implicates that multiple functional brain networks support active maintenance of the attended memory objects. Neuroimaging research across various modalities suggests that both domain-general networks related to attention (i.e., fronto-parietal network) and cognitive control (i.e., cingulo-opercular network), and the sensory-specific neural regions are involved in maintaining items held in working memory (Goldman-Rakic, 1995; Harrison and Tong, 2009; Higo et al., 2011; Ester et al., 2015). For example, auditory and verbal working memory research has demonstrated contributions of the domain-general networks, language-related brain regions (e.g., the inferior frontal gyrus and inferior parietal areas), and premotor cortex (Baddeley, 2003; Hickok and Poeppel, 2007; Koelsch et al., 2009; Fedorenko et al., 2011; Buchsbaum et al., 2005), as well as the bilateral auditory cortices for retaining stimulus-specific features of auditory information in memory (Linke et al., 2011; Linke and Cusack, 2015; Kumar et al., 2016).

Collectively, the converging evidence suggests that neural traces of memory representations under focused attention are maintained across distributed brain networks.

However, less is known about whether and how these multiple networks support the benefits from attention re-directed to working memory. Especially, while selective attention is critical in forming and maintaining stable auditory objects in memory (Fritz et al., 2007; Shinn- Cunningham, 2008; Wilsch and Obleser, 2016), how selective attention-to-memory affects the internal representations of auditory working memory and the associated neural implementations is not well specified.

Prior auditory working memory studies demonstrate that both featural and object- based attention re-directed to memory representations guided by retro-cues improves memory performances (e.g., Kumar et al., 2013; Backer and Alain, 2014; Backer et al., 2015; Lim et al., 2015, 2018; see Backer and Alain, 2012; Zimmermann et al., 2016 for reviews). In particular, studies utilizing psychophysical modeling demonstrate that retrospective auditory attention enhances representational precision of the attended versus unattended auditory objects held in memory (Kumar et al., 2013; Lim et al., 2015, 2018). Importantly, such precision benefit has been reflected in an attention-induced increase in neural activities measured via EEG (i.e., enhanced sustained negativity and neural alpha (∼10 Hz) oscillatory power), indicating that greater cognitive resource is allocated to the attended memory object. This work supports the notion that retrospective auditory attention enhances the precision of the memory item, rather than reducing cognitive load by removing the unattended items from memory (Lim et al., 2015); however, the limited spatial resolution of EEG cannot delineate how distinct functional networks of the brain contribute to precision benefit from retrospective attention.

A few existing fMRI studies provide consistent evidence that retrospective attention to auditory memory representations actively engages fronto-parietal cortical regions as well as modality-specific auditory cortex. This evidence suggests that orienting attention to, and selective accessing of, auditory memory representations rely upon both top-down control of attention and perceptual processes (Buchsbaum et al., 2005; Johnson et al., 2005; Backer et al., 2020). Nevertheless, how these distinct regions across the brain—cognitive control vs. auditory processing regions—support memory representational enhancement via retrospective attention remains unclear.

To investigate the neural underpinning of auditory retrospective attentional benefit, we conducted an fMRI experiment, in which participants performed a previously established auditory working memory task—a delayed recall of syllable pitch discrimination task (Lim et al., 2015; Alavash et al., 2018; Lim et al., 2018). In this task, listeners encoded two speech syllable sounds into memory and then received a retroactive cue to direct attention to one of the syllables (i.e., auditory objects) prior to recalling lower-level feature (pitch) of the object. In order to specifically model the parameter for precision at which each participant recalled the syllable object from memory, we probed fine-grained parametric changes of the low-level feature (e.g., Bays & Husain, 2008; Zhang & Luck, 2008; Murray et al., 2013; Lim et al., 2015, 2018). Here, we specifically investigated (*i*) whether selective attention to auditory memory enhances representational fidelity by using a psychophysical modeling approach (Bays and Husain, 2008; Zhang and Luck, 2008; Murray et al., 2013); (*ii*) how retrospective attention modulates the neural activities in domain-general and modality-specific brain regions by using univariate fMRI analysis; and (*iii*) whether the speech memory objects were retained in aforementioned brain regions, and which of these brain regions contribute to the attentional enhancement in mnemonic fidelity using a multivariate classifier approach on fMRI data.

We hypothesized that if retrospective attention facilitates working memory performance by enhancing mnemonic fidelity of the attended object, we would observe higher precision in recalling the acoustic feature of the syllable object held in memory, and attention-related increase in neural activations in the domain-general brain regions, reflecting the higher attentional demands to maintain precise memory representations. In addition, we expect that the auditory memory object representations distributed across the brain—the domain-general cognitive control and sensory-specific brain regions—would contribute to the attentional enhancement in mnemonic fidelity, but potentially in varying degrees given that functionally discrete brain regions may retain either more abstracted or sensory-specific information in working memory (Lee et al., 2013; Ester et al., 2015; Christophel et al., 2017). However, if retrospective attention reduces working memory load by removing unattended items from memory (Kuo et al., 2012; Souza et al., 2014), we expect to observe an attention-related decrease in neural activations in the domain-general networks (i.e., fronto-parietal and cingulo-opercular networks), and no direct relationship between the neural representations of the auditory memory object to the mnemonic precision in recalling the object.

## Materials and Methods

### Participants

Twenty-two native German speakers (mean age: 27.9 years; SD: 2.75 years; age range: 25– 35; 12 females) participated in the experiment that has been a part of a previously published study (Alavash et al., 2018). Participants were recruited from the participant database of the Max Planck Institute for Human Cognitive and Brain Sciences (Leipzig, Germany). Two additional participants underwent the study, but were excluded from data analyses due to excessive head movements (>5 mm). All participants reported normal hearing, and no history of neurological or psychiatric disorders. None of the participants were under medication. All participants except one were right handed. Based on our prior EEG and behavioral work adopting the similar auditory retro-cueing task (Lim et al., 2015, 2018), we determined that with 17 participants we would achieve at least 80% power to detect the within-subject effect of retro-cues even under simultaneous noise in the background.

All participants gave their informed written consent, and received financial compensation for their participation. The study procedure was in accordance with the Declaration of Helsinki. The local ethics committee (University of Leipzig) approved the study.

### Stimuli

The stimulus material was identical to the ones used in previously published studies (Alavash et al., 2018; Lim et al., 2015, 2018). In brief, we used two German syllables, /da/ and /ge/. Each syllable category consisted of six naturally spoken tokens, truncated from German words recorded by a trained female speaker. All sound tokens were digitized at 44.1 kHz, and 200- ms in duration. For each token, 3-ms and 30-ms linear ramps were applied at sound onset and offset, respectively.

For each syllable category, two tokens served as to-be-probed syllables in the syllable- pitch discrimination task. The pitch dimension (i.e., the fundamental frequency; F0) of each token was parametrically manipulated in four steps: ±0.125 and ±0.75 semitones from its F0; we chose these relatively small step sizes in order to closely examine whether selective attention to speech objects held in memory benefits precise recall of low-level (pitch) information of the cued object. The original token was presented during the syllable encoding phase, and the F0-manipulated tokens were presented as probes. The average F0’s of the to-be-probed /da/ tokens were 162.8 Hz [min = 154.0, max = 171.7] and 175.5 Hz [min = 169.4, max = 183.5] for /ge/ tokens. To increase acoustic variability during the syllable encoding phase, we created additional 12 tokens for each syllable category based on the six utterances from the same speaker. These sounds served as unprobed tokens: they were presented during encoding, but were not used for discriminating pitch change at probe. On average, unprobed tokens of /da/ and /ge/ had F0’s of 163.7 Hz [min= 152.8, max = 181.7] and 175.9 Hz [min = 163.7, max = 186.9], respectively. These unprobed sounds were used to ensure that participants were exposed to acoustically-variable sounds beyond the fixed set of to-be-probed tokens, but within the similar F0-range such that to-be-probed and unprobed sounds were indistinguishable during the encoding phase.

All stimuli were normalized to equivalent amplitude (root-mean-squared dB full scale; RMS dBFS). Praat ver. 5.3 was used to manipulate the F0 dimension.

### Experimental Design

In the MRI scanner, participants performed a previously established auditory working memory task—a syllable-pitch discrimination task with retroactive cues (Alavash et al., 2018; Lim et al., 2015, 2018). Figure 1A illustrates the task structure. On each trial, participants first heard two syllable tokens (i.e., /da/ and /ge/) in a random order sequence, separated by a 1- or 2-s inter-stimulus interval (ISI). After a delay period of 3–5 s, participants saw a visual retro-cue for 1 s displayed on the screen. With an additional delay of 5–7 s following the visual cue, participants heard a probe syllable. The probe was one of the two syllables heard during encoding with a slight manipulation in the pitch (F0). Within a 4-s time window, participants responded whether the pitch of the probe syllable was higher or lower than the corresponding syllable (i.e., the same category) heard during encoding in the beginning of the trial. Response feedback was provided on the screen for 0.5 s. Trials were separated by a 3–7 s silence period with a fixation cross on the screen.

**Fig. 1.**
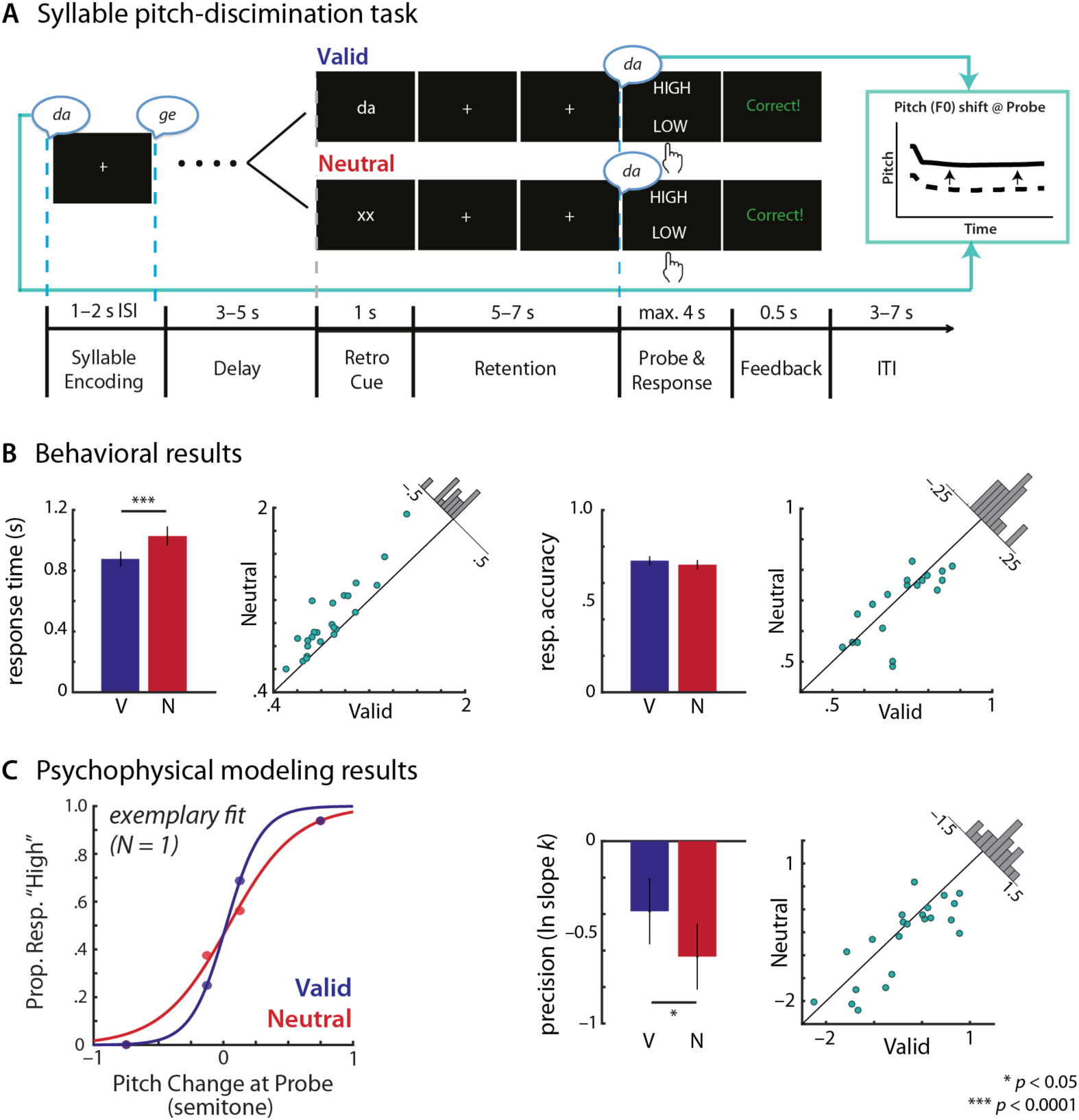
Illustration of the syllable pitch-discrimination task and the summary of task performance and psychophysical modeling results for validly cued (V) and neutrally cued (N) trials. (A) Trial structure of the task. On each trial, participants encoded two syllables (each syllable 0.2 s in duration) presented in a random order separated by a 1- or 2-s inter-stimulus interval (ISI). After the first delay period (3–5 s), a visual retro-cue appeared on the screen for 1 s, which was followed by a stimulus-free retention period (with duration jittered from 5–7 s). After the retention period, participants heard one of the encoded syllables again as a probe, the pitch of which was slightly manipulated. Participants judged whether the pitch of the probe syllable was higher or lower than the corresponding syllable heard during encoding (i.e., beginning of the trial). Participants were given a maximum of 4 s to provide their response. On every trial, participants received either a valid or neutral retro-cue, which respectively informed or did not inform the identity of the to-be-probed syllable category. (B) The behavioral performance results as a function of retro-cues. Left, response time of the correctly responded trials. Right, response accuracy. Bar plot shows the average performance measure in the valid and neutral cue conditions. Error bars indicate ± 1 standard error of the mean (SEM). Scatter dots represent the corresponding behavioral measure with respect to the cue conditions of individual participants. The 45-degree lines represent the performances in the two retro-cue conditions are identical. The histograms represent the distributions of the cue-related behavioral difference across N=22 participants. (C) The parameter estimates of psychophysical modeling results. Left, an exemplary psychophysical modeling fit of a single participant’s response patterns in the valid and neutral retro-cue conditions. Each dot indicates the proportion responses “high” as a function of syllable pitch change at probe, with respect to the matching reference syllable presented during the encoding phase. The lines denote the logistic model fits. Right, individual’s perceptual precision estimates (ln slope *k*) with respect to the cue conditions. The illustration scheme is identical to panel B.

During the retention period on each trial, participants saw one of the two retro-cues: valid or neutral. A valid retro-cue indicated the identity of the probe syllable category; a written syllable (e.g., “da”) was presented on the screen for 1 s in order to direct participants’ attention to one of the syllables held in memory prior to hearing the probe. On the neutral retro-cue trials, participants were presented with “xx” on the screen. This cue did not provide any information about the upcoming probe syllable. Thus, listeners must maintain both syllable sounds in memory until hearing a probe.

The pitch variations of the two to-be-encoded syllables were drawn from the similar F0-range. On average, pitch (F0) difference between the two syllables presented during encoding was 4.13 Hz (cf. note that a wider pitch range (>200 Hz) of auditory pure tones were used in previous auditory working memory studies; e.g., Link et al., 2011; Kumar et al., 2016), and the F0 difference was not significantly different between the two retro-cue conditions (*p* = 0.998). Nevertheless, F0’s of the syllables presented during the encoding phase varied continuously across trials; thus, listeners must remember specific sound instance of the syllable in order to perform the task.

Participants performed a total of eight runs of 16 trials. On each run, the valid and neutral retro-cue trials appeared on an equal probability (i.e., 8 trials per cue condition); thus, participants were unaware of trial type until seeing a visual retro-cue. For every two runs, unique combinations of 2 syllable categories × 2 syllable positions during encoding being probed × 2 retro-cue types × 4 probed pitch step sizes were presented.

### Procedure

Prior to fMRI scanning, all participants completed a short practice session of the syllable pitch discrimination task on a laptop in a separate behavioral testing room. The practice session involved 18 trials (9 trials for each cue type), which probed syllable pitch changes that varied in ±1.25 semitones. This practice session was given to ensure that participants understood the task prior to scanning. We ensured that all participants were able to perform the task >80% on this practice session.

In the scanner, participants used left and right index fingers to provide “high” and “low” responses. The mapping between the responses and hands were counterbalanced across participants. Experimental trials were controlled with Presentation software (Neurobehavioral Systems). Auditory stimuli were delivered through MR-compatible headphones (MrConfon, Magdeburg, Germany), and Music safe pro earplugs (Alpine Hearing Protection) provided an additional attenuation (on average 20-dB across 125–8000 Hz range). To ensure that participants could hear the auditory syllables during task in the scanner, we conducted a brief hearing test; participants heard four randomly selected speech stimuli with an on-going functional (T2*-sequence) scanner noise, and verbally reported the syllable categories of the sounds.

After the eight scanning runs of the syllable-pitch discrimination task, participants went through two additional scanning runs of an auditory habituation (∼16 min), involving listening to a sequence of four syllable tokens while watching a silent movie. However, excessive head movements were observed in at least 7/22 participants (> 4.5 mm) during this task; therefore, data from this task was not further reported.

Note that the data and results reported here have been acquired as part of a larger, pharmacological fMRI study such that all participants went through two identical fMRI sessions, separated by at least one week (EudraCT number 2015-002761-33). The study design was a double-blind, counterbalanced repeated-measures of the intake of levodopa (L-dopa) and placebo. Because our current research focus is on the neural basis of retrospective attention to auditory working memory (rather than the effect of L-dopa), we report the results only from the placebo session. Of significant note, in all our analyses, we controlled a potential confound in the placebo session order differences across participants. The effect of L-dopa on auditory working memory, corresponding functional neural organization, and the detailed procedure of the pharmacological fMRI study has been described elsewhere and can be found in Alavash et al. (2018).

### Statistical Analysis

#### Behavioral measures and psychophysical modeling analysis

Response time (RT) of the correct trials and accurate syllable pitch judgment (i.e., a binary response accuracy measure) served as overall behavioral performance measures. To ensure normality, RT data were log-transformed prior to analysis. These single-trial behavioral measures were separately analyzed using a (generalized) linear mixed-effects model framework (*lme4*, ver. 3.3) in R environment. The model included a fixed factor of retro-cue (Valid vs. Neutral), which was the main variable of interest. The model also included the session order as a nuisance factor. The random-effects terms included by-participant intercepts. The significance of the effects was determined based on Wald χ^2^ tests (at *p* < 0.05). Beyond these two overall behavioral performance measures, we obtained a more fine-grained perceptual precision measure for each participant and for each condition using a psychophysical modeling approach. We adopted this approach as it is commonly used in psychophysics research spanning across various domains (e.g., Bays and Husain, 2008; Zhang and Luck, 2008; Murray et al., 2013; Kumar et al., 2013; Rohenkohl et al., 2012; Myers et al., 2014; Herbst and Obleser, 2019). We estimated individual participant’s perceptual precision in judging syllable pitch change by fitting each participant’s response patterns across the parametric changes in the pitch of the probe syllable relative to that of the encoded syllable. Following our previous auditory working memory work (Lim et al., 2015, 2018), we used a nonlinear least square curve fitting procedure (MATLAB *lsqcurvefit*) to model individual’s response patterns with a logistic (i.e., psychometric) function,

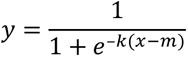

where *x* indicates the amount of pitch changes occurred at probe relative to the encoded syllable (4 step sizes), *y* indicates the proportion of “high” responses, *k* indicates the slope of the curve, and *m* indicates the mid-point/inflection point of the function along the x-axis. The parameter *m* estimates bias in each participant’s response, and the slope *k* provides an estimate of the perceptual precision in pitch judgment in the task: higher perceptual precision is indicated by steeper slope of the curve (see Fig. 1C, left for a single participant exemplary model fit).^1^ As done for analyses of the behavioral measures, the psychophysical modeling estimates of perceptual precision slope *k* and bias *m* were separately analyzed using a linear mixed-effects model. The linear mixed-effects models included a fixed factor of retro-cue (Valid vs. Neutral) and the nuisance factor of the session order. The random-effects term included by-participant intercepts. The significance of the effects was determined based on Wald χ^2^ tests (at *p* < 0.05).

#### Functional MRI acquisition and analysis

Imaging data were acquired on a 3-Tesla Siemens MAGNETOM Prisma^fit^ scanner (TimTrio Upgrade) at the Max Planck Institute for Human Cognitive and Brain Sciences. Whole-brain functional images, sensitive to the blood-oxygen-level-dependent (BOLD) contrast, were acquired with a 20-channel head/neck coil using a T2*-weighted echo-planar imaging (EPI) sequence (repetition time (TR) = 2000 ms; echo time (TE) = 26 ms; flip angle (FA) = 90°). Each volume had 40 oblique axial slices (ascending) parallel to the anterior commissure– posterior commissure (AC–PC) line (acquisition matrix = 64 × 64; field of view (FOV) = 192 mm × 192 mm; voxel size = 3 × 3 × 3 mm; interslice gap = 0.3 mm). For each block of the syllable-pitch discrimination task runs, a total of 181 volumes were acquired.

Structural images of (*n* = 15) participants were available from the Institute’s brain database acquired from previous MR studies. A magnetization prepared rapid gradient echo (MP-RAGE) T1-weighted sequence was used to acquire the structural images (TR = 2300 ms; TE = 2.01–2.98 ms; FA = 9°; 176 saggital slices with 1 × 1 × 1 mm resolution). For the participants (*n* = 7) whose structural images did not exist in the database, a high-resolution T1-weighted structural image was acquired using a MP-RAGE sequence (TR = 2300 ms; TE = 2.98 ms; FA = 9°; 176 saggital slices with 1 × 1 × 1 mm resolution) at the end of functional scans.

Imaging data were preprocessed using the Analysis of Functional NeuroImages (AFNI; ver. 01-2017) software (Cox, 1996). The first two volumes of each run were discarded to allow the scanner to approach net magnetization equilibrium. Preprocessing steps included 1) slice-time correction, 2) motion correction of all volumes based on the first volume of the syllable- pitch discrimination task run using a rigid-body alignment, 3) spatial alignment of functional data to each participant’s structural anatomy, and 4) the signal for each voxel was scaled to a global mean of 100. For univariate analysis, images were spatially smoothed with an 8-mm full-width at half maximum (FWHM) Gaussian kernel. For multivariate classification analysis, data were not smoothed. Spatial normalization of functional images to Montreal Neurological Images (MNI–152) was performed prior to the group-level statistical analyses.

##### Univariate analysis

For each participant, a voxel-wise general linear model (GLM) was constructed to estimate hemodynamic responses during different phases of the syllable- pitch discrimination task across all scanning runs. The BOLD response to the trial onset (i.e., the syllable encoding phase) was modeled using a canonical gamma variate function (GAM by AFNI) along with a pairwise amplitude-modulated regressor that accounted for the ISI between presentation of the two syllables. The second task regressor captured the entire hemodynamic response of the visual retro-cue and the following stimulus-free retention phase up to auditory probe phase using a finite impulse response (FIR) function (TENT by AFNI) as a change in BOLD activation across 0–20 s from the retro-cue onset. To account for the varying durations of the retention phase (Fig. 1A), a parametric amplitude-modulated regressor was entered as a trial-wise covariate. The third regressor captured the BOLD response related to the participants’ response and visual feedback, we used a canonical GAM. All task regressors were separately modelled for the valid and neutral retro-cue trials in the GLM. The baseline activity was modelled by trends from linear up to fourth-degree polynomials in order to remove the slow scanner signal drift (i.e., high-pass filter with an approximate cut-off of 240 s). The six additional continuous head motion parameters were entered as regressors of no interest.

The primary purpose of the univariate analysis was to identify brain regions that were differentially modulated by selective vs. non-selective attention directed to auditory working memory objects informed by retro-cues. Thus, we conducted a group-level analysis to contrast the neural activities during the cue and the following retention phase in the valid vs. neutral retro-cue conditions. For each participant, the cue-retention phase neural activity was quantified by averaging the *β*-parameter estimate of the FIR during 4–8 s from the cue onset, separately for the valid and neutral retro-cue conditions (Fig. 2B). The group-level analysis was conducted using a voxel-wise linear mixed-effects modeling approach (3dLME by AFNI). The model involved one within-subjects fixed factor of interest (retro-cue: Valid vs. Neutral) with participants as a random factor, and regressor of no interest (i.e., placebo session order) was centered, and entered as covariates in the model. Planned contrast was established to compare BOLD activations during the cue–retention phase in the valid vs. neutral retro-cue conditions. The group-level maps were corrected for multiple comparisons at the voxel-level using false discovery rate (FDR *p* < 0.05; Benjamini and Hochberg, 1995).

**Fig. 2.**
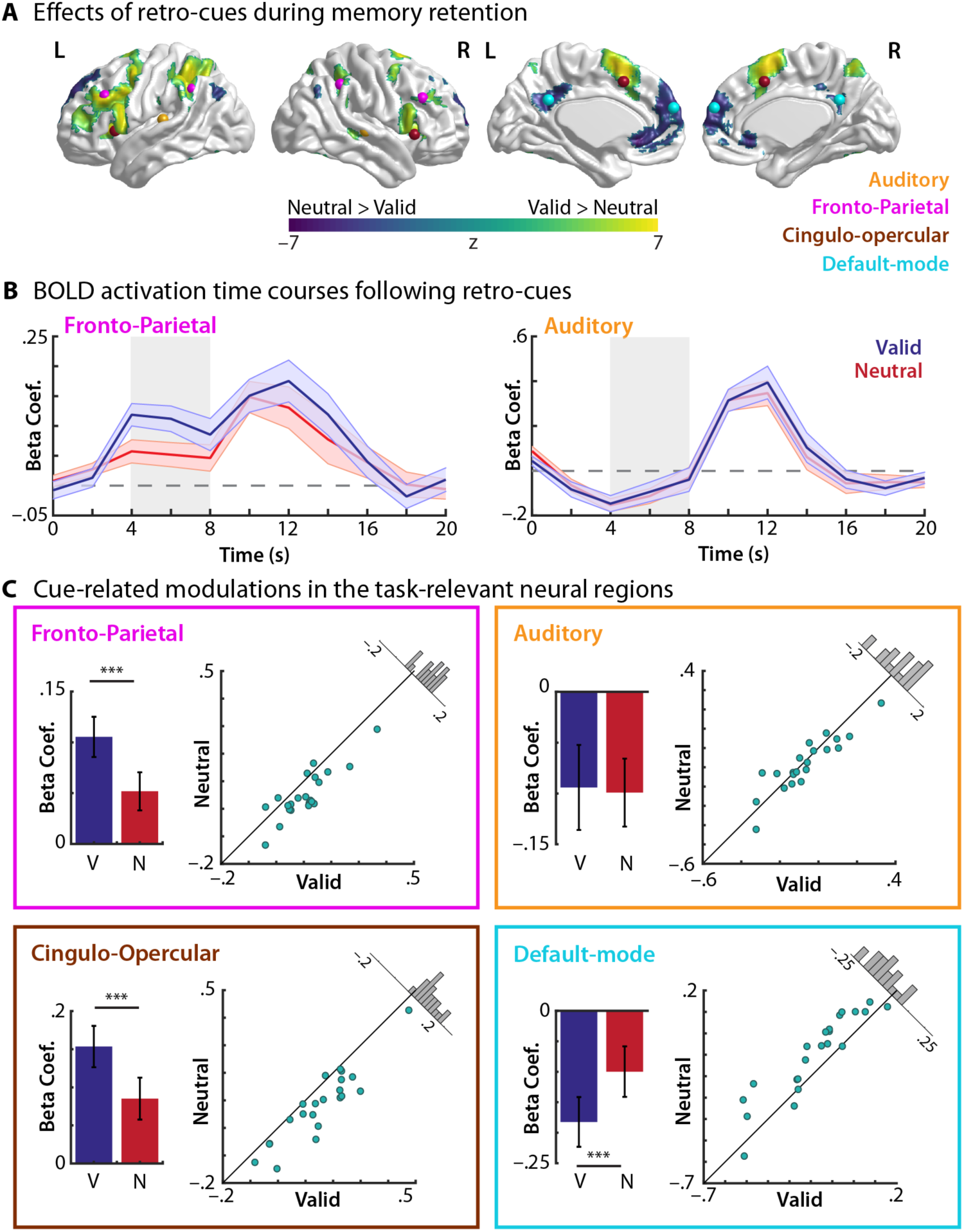
Result of the univariate analysis of Valid vs. Neutral retro-cues on modulating the brain activation during the post-cue memory retention phase. (A) Brain regions exhibiting significant effects of retro-cues (Valid, V – Neutral, N) (FDR corrected at *p* < 0.05). ROI seeds of the four, independently-defined networks are illustrated with the spheres of the corresponding colors. L, left; R, right. (B) BOLD activation time courses of the Valid vs. Neutral retro-cue conditions from the cue onset. The BOLD time courses were extracted from the exemplary regions (fronto-parietal and auditory ROIs). The grey boxes denote the time windows of the BOLD activity during the cue and the following memory retention phase (4–8s in FIR; see Methods and Materials). (C) Cue-related modulation of the BOLD activation in each network ROI. Bar graphs depict the average BOLD activations (i.e., beta coefficient) during the time points (4–8 s) of the Valid and Neutral retro-cue conditions. Scatter plots represent the average BOLD activations of the corresponding ROIs with respect to the retro-cue conditions of individual participants. The 45- degree lines indicate identical magnitude of activation of the two cue conditions. The histograms depict the distributions of the cue-related BOLD activity differences across N=22 participants. Error bars in panels B and C indicate ± 1 SEM. *** indicates *p* < 0.0001 (FDR-corrected).

##### Region-of-interest (ROI)-based analysis

We conducted an additional ROI-based analysis (*i*) to complement and confirm the whole-brain univariate analysis as to whether the valid vs. neutral retro-cues modulate the brain activities in the well-known domain-general cognitive networks; and, more importantly, (*ii*) to investigate whether the neural activity modulations in these distinct networks are related to behavior (see below). To ensure independence of defining the domain-general network ROIs and the retro-cue-related BOLD activity modulations, we defined the ROIs based on the brain regions organized by intrinsic connectivity networks established in previous literature (Dosenbach et al., 2007; 2008; Power et al., 2011; Sadaghiani and D’Esposito, 2014). To this end, we placed spherical seeds (radius 6 mm) at the centers of each brain network coordinates defined in previous work (Fig. 2A); for the fronto-parietal network, the seeds were placed over the bilateral dorsolateral prefrontal cortex (MNI coordinates: [–44, 27, 33] and [46, 28, 31]) and the bilateral inferior parietal lobules ([-53 –50 39], and [54 –44 43]) as defined in Dosenbach et al. (2007) and Power et al. (2011). ROIs representing the cingulo-opercular network were placed based on Dosenbach et al. (2007) in the bilateral insula/frontal operculum ([–36, 18, 2] and [38, 21, –1]), and the supplementary motor area (SMA)/dorsal anterior cingulate cortex at [0, 15, 45]. Default mode network seeds were placed at the posterior cingulate cortex [1, –51, 29] and medial prefrontal cortex [–1, 61, 22] based on previous findings (Raichle et al., 2001). In order to observe neural activity modulated by the retro-cues in the sensory processing (i.e., auditory) regions, we also placed spherical seeds, centered over the peak activations during encoding of the two syllables at the onset of trial in the current experiment; these were placed in the bilateral auditory regions (superior temporal gyrus/sulcus) at [–58, –24, 12] and [62, –22, 2].

To test whether and how the retro-cues modulate brain activity in each of the four network ROIs, we first extracted the average BOLD responses during the cue–memory retention phase (i.e., average *β*-estimates during 4–8 s time window of the FIR; Fig. 2B) from the voxels within the seeds representing each network. The average *β* estimate data of each network were submitted to a linear mixed-effects model with a fixed factor of retro-cue (Valid vs. Neutral), a nuisance regressor of the session order, and by-participant random intercept. We corrected for multiple comparisons using FDR (*p* < 0.05).

##### Multivariate analysis

Our main goal was to understand how object-based attention directed to auditory working memory improves the representations held in memory. To this end, we conducted a multi-voxel pattern analysis (MVPA) to examine (*i*) whether the attended syllable object information held in memory can be decoded from patterns of neural activity during the memory retention phase, and (*ii*) whether the decoding performance for the attended auditory object differed between the valid vs. neutral retro-cue conditions. Here, we performed a whole-brain searchlight analysis with a searchlight sphere of 10 mm radius (∼123 voxels; Kriegeskorte et al., 2006; Haynes et al., 2007). For each retro-cue condition, we used a linear support vector machine classifier, implemented in LIBSVM (Chang and Lin, 2011), with neural activation estimates in voxels in a searchlight sphere as features.

Two syllable sounds presented during encoding varied throughout the experiment, and they were not so differentiated by the pitch dimension (F0 difference varying between 0.64–7.86 Hz across trials), we trained the classifiers to decode syllable category information (i.e., /da/ vs. /ge/) instead decoding the syllable pitch. To identify brain regions that retain auditory-based form of the attended syllable category during memory retention, we trained SVM classifiers to decode syllable categories from the brain activity patterns when participants heard one of the two syllables at the end of the trial (i.e., the auditory probe phase) so that neural activity patterns used for training was not contaminated by other auditory events. We then tested whether the trained classifier could successfully decode syllable category information from the brain activity patterns during the memory maintenance phase (i.e., the retro-cue and the following stimulus-free retention period). Of important note, it is possible that in valid retro-cue trials, neural activity evoked by auditory probes might exhibit syllable category-specific bias informed by the valid cues. To avoid this potential bias, classifier training was performed only on the auditory probe phase of the neutral retro-cue trials, in which participants did not have any expectation about the upcoming probe syllable. Thus, training and testing phases for the classifier were separate and temporally independent from each other (especially, for the valid retro-cue trials).

To estimate neural activity patterns during the cue-retention and the auditory probe phases used for the MVPA, we first constructed a single subject-level GLM for each functional run. As in the univariate analysis, a 0–20 s FIR was used to capture the hemodynamic responses ranging from the retro-cue onset, the following retention, to the auditory probe phases of the task. Four, 20-s FIRs were included in the GLM to separately model the hemodynamic response of each condition: 2 cue types (valid and neutral) × 2 to-be-probed syllable categories (i.e., /da/ and /ge/). From each run-wise GLM, condition-specific activation across voxels were estimated with *t*-estimates for the MVPA (Misaki et al., 2010). The neural activity during the cue-retention phase was quantified by averaging the *t*-estimates of the FIR during 4–8 s from the cue onset; and the neural activity evoked by the auditory probe was quantified by averaging the *t*-estimates during 10–12 s of the FIR. These time windows were selected to account for about 4–6 s delay of the hemodynamic responses to these trial events (see Fig. 2B for the average activations during all time points of the FIR). The GLM also included additional canonical regressors to account for BOLD responses during encoding of two speech syllables during trial onset, and during participants’ response and visual feedback, as well as continuous regressors to model head motion and scanner drift.

For cross-validation of the generalization of classifier performance, eight functional runs were divided into training and test datasets. The classifier was trained on the auditory probe phase activity of the neutral-cue trials of five randomly selected functional runs, and the classifier was tested on the cue–retention phase activity of each cue condition from the remaining three functional runs. The cross-validation iteration was performed 28 times (i.e., a half of all possible combinations of five training and three test datasets), and the decoding accuracy was calculated based on the average performances across all iterations, separately for each cue condition. Searchlight MVPA was performed on unsmoothed functional data in each participant’s native space. The resulting decoding accuracy map of each participant was centered at zero (i.e., accuracy minus 50% chancel level), and normalized to standard MNI space. The group-level accuracy map was determined by averaging the classification accuracy across participants.

To determine statistical assessment of classification performance above chance, we adopted a non-parametric permutation approach in order (Stelzer et al., 2013; Allefeld et al., 2016). To this end, we repeated the whole-brain searchlight analysis with randomly assigned classification labels for 1000 times for each participant. We then generated a permuted group-level accuracy map by averaging one randomly selected map out of the 1000 permuted maps drawn from each participant (with replacement); this process was repeated 10^5^ times, which resulted in 10^5^ permuted group-level accuracy maps. Based on the null-hypothesis distribution (permutation) maps of chance level decoding, we computed a map of voxel-wise significance value for the un-permuted group accuracy map, and corrected for multiple comparisons (FDR at *p* < 0.05). From these regions that survived the statistical threshold, we extracted decoding accuracies for the valid and neutral retro-cue conditions.

##### Brain–Behavior relationships

In order to understand how the network of task-relevant brain regions contribute to the representational precision benefits from valid retro-cues (i.e., Valid – Neutral: Δ ln slope *k*), we performed a multiple linear regression analysis to examine the relationship between the degrees of neural modulations and the representational precision modulations from the retro-cues (i.e., Valid vs. Neutral) across individuals. The two types of neural modulation measures, one from the univariate and the other from the multivariate results, were analyzed separately to explain the extent of the precision enhancement with valid (vs. neutral) retro-cues.

Starting from a model that includes all factors representing neural modulations of multiple brain regions, we used a step-wise model comparison to find the best model that explains the precision modulations from valid vs. neutral cues. The model comparison was based on the AIC of the model (*StepAIC* from the package *MASS* ver. 7.3-49 in R) as well as *χ*^2^ testing to deduce into a simplest model that explains the cue-related modulations of the perceptual precision.

In analyzing the univariate BOLD data, we performed the model comparison, starting from the full model—that is, the neural activity modulations derived from the four ROIs, including the bilateral auditory region (Fig. 2), to explain the degree of the perceptual precision modulations (Δ ln slope *k*):

Δ ln slope *k* = Δ Auditory + Δ Fronto-Parietal + Δ Cingulo-Opercular + Δ Default-mode

where Δ represents the degree of cue-related modulation (i.e., Valid – Neutral).

Using a similar approach, we analyzed the multivariate data to understand whether the neural decoding accuracy modulations in the task-relevant brain regions were related to the extent of precision modulation from valid vs. neutral cues. To this end, we started the model comparison from the task-relevant cortical regions that exhibited significantly accurate decoding of to-be-probed syllable categories during the memory retention period (Fig. 3; Table 2). These regions were selected based on whether they fell within the language processing regions (Baddeley, 2003; Hickok and Poeppel, 2007; Koelsch et al., 2009), auditory regions in the temporal cortex, domain-general attentional and cognitive control network regions (i.e., fronto-parietal and cingulo-opercular networks). Starting from the full-factor model that included the neural decoding accuracy modulations from 13 brain regions (Table 2), we used a step-wise model comparison to deduce from the full-factor model to find the simplest/best model that explains the extent of precision modulation (Δ ln slope *k*: ln slope *k*Valid – ln slope *k*Neutral).

**Fig. 3.**
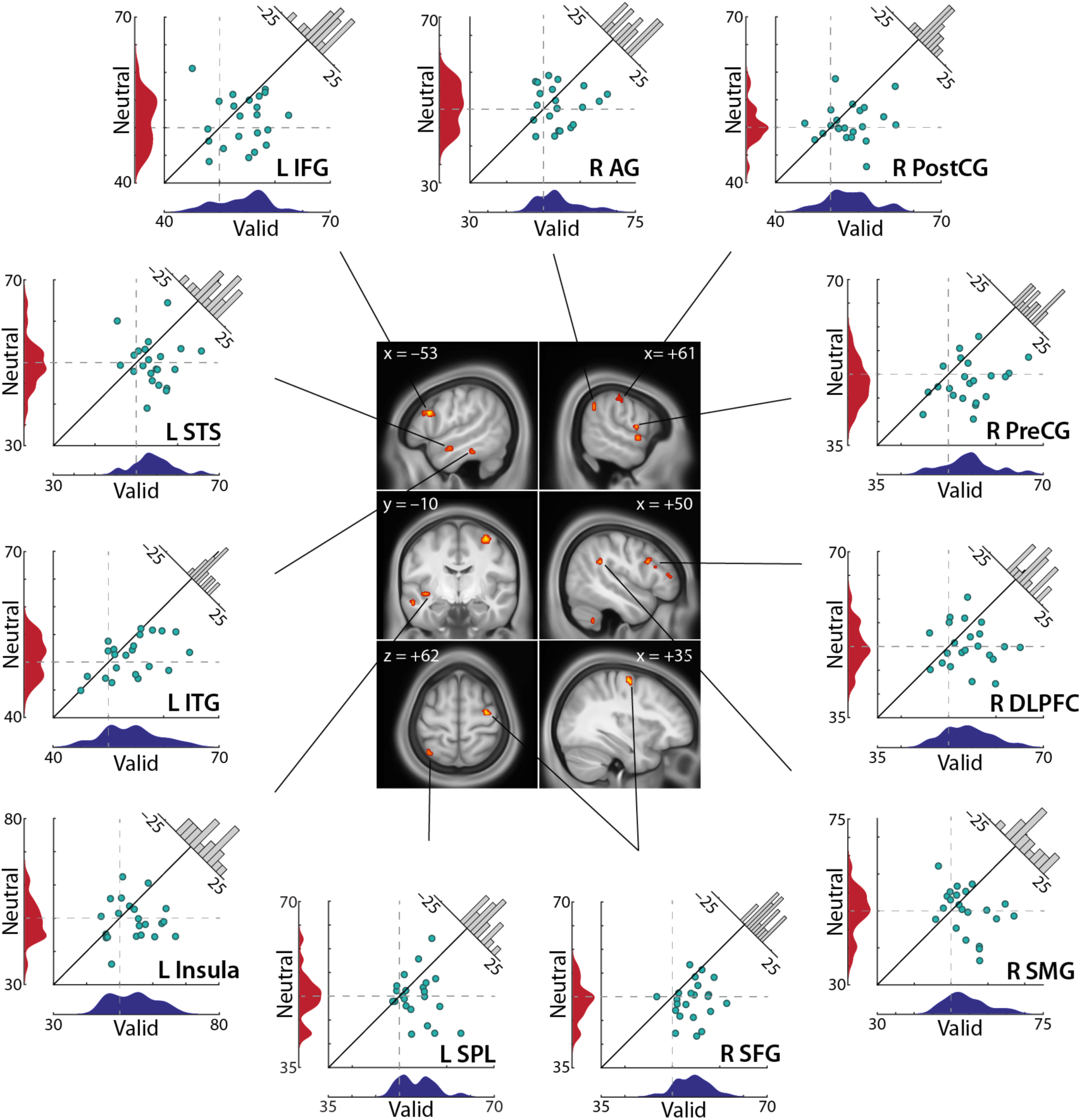
Multivariate classification results on decoding attended syllable object held in memory. The classifier was trained to decode syllable categories based on the auditorily presented probe sounds in the neutral retro-cue trials; and the classifier was tested to decode the neural activity patterns during the memory retention phase following retro-cues. The brain regions exhibit above-chance classifier accuracies on decoding the cued syllable category (i.e., indicated by valid retro-cues; see Methods and Materials for further details) against non-parametric testing. Each scatter plot depicts individuals’ decoding accuracies of the valid vs. neutral retro-cue conditions extracted from the corresponding brain regions that retain neural representations of syllable category. The distribution of decoding accuracy of each cue condition is shown next to the corresponding axis of the scatter plot. The histogram represents the distributions of the cue-related decoding accuracy modulations (Valid – Neural) of N=22 participants. The 45-degree line would denote identical decoding accuracies in the two cue conditions. Dashed lines are shown to indicate hypothetical chance level (50%) decoding accuracy.

Note that all of the cue-related modulation data (i.e., the perceptual precision modulation and neural modulations across the brain regions) were z-scored prior to the model testing. In order to ensure the validity of the final model resulted from the model comparison step, we performed a leave-one-participant-out cross-validation (i.e., 22 CV’s). On each CV step, we used *n*=21 participants’ data to fit the model parameters, and tested on the remaining one participant’s data. We compared the cross-validated fitted values vs. the actual data outcome using a correlation test; and we report the average adjusted R^2^ from 22 cross-validation steps.

## Results

### Valid retro-cues enhance syllable-pitch memory recall speed and precision

We first examined the effect of retro-cues (valid vs. neutral) on behavioral performances. A linear mixed effects model on participants’ log-transformed response times (RTs) of the correct trials revealed a significant main effect of retro-cues (MValid–Neutral = –151.029 ms, *χ*^2^(1) = 92.264, *p* < 0.0001). As shown in Fig. 1B (left), participants were significantly faster in correctly judging the pitch of syllables when they received valid retro-cues compared to the neutral cues. On the contrary, we did not observe robust benefit from valid retro-cues on response accuracy. Although we observed 13/22 participants were more accurate in recalling syllable pitch with valid than neutral retro-cues, a logistic mixed-effects model on listeners’ response accuracy did not reveal any significant effect of retro-cues (*χ*^2^(1) = 1.729, *p* = 0.189; Fig. 1B, right).

We further examined the effect of retro-cues on listeners’ perceptual precision and response bias, estimated by the slope (*k*) and inflection/mid-point (*m*) parameters of logistic function fit from psychophysical modeling (see Materials and Methods; Fig. 1C, left). A linear mixed-effects model on the log-transformed perceptual precision estimate (ln slope *k*) revealed a significant effect of retro-cue condition (MValid–Neutral = 0.248, *χ*^2^(1) = 4.562, *p* = 0.0327). Consistent with the prior findings (Lim et al., 2015; 2018), valid retro-cues led to more precise recall of syllable pitch compared to the neutral retro-cue trials (Fig. 1C, right). For the response bias estimate (*m*), a linear mixed-effects model revealed no significant effect of retro-cue condition (*χ*^2^(1) = 1.216, *p* = 0.270); moreover, neither of the cue conditions did exhibit significant bias (one sample *t*-test against 0, Valid: *t*(21) = –0.433; *p* = 0.670; Neutral: *t*(21) = 1.112, *p* = 0.279).

Note that for any of these measures, we did not find any significant main or interaction effects related to the factor of no interest—that is, the order of the two separate scanning sessions taking place for each participant (all *χ*^2^s < 0.359, all *p*s > 0.549). In addition, we performed the same analyses with an additional factor to test whether the probed syllable category had any impact on these measures; except for the response accuracy exhibiting a slightly higher accuracy for /da/ than /ge/ (*χ*^2^(1) = 5.028, *p* = 0.020; M ± SD = 3.906 ± 6.900%), we did not find any significant effects related to the probed syllable category in any other measures (all *p*s > 0.136).

### Valid and neutral retro-cues differentially modulate domain-general networks during memory maintenance

As shown in Fig. 2A, the univariate analysis revealed significantly greater recruitment of broad regions across the whole brain during memory retention when participants received valid than neutral retro-cues (Table 1). While the effect was more prevalent in the left hemisphere, these regions include the bilateral inferior frontal gyri extending into the dorsolateral prefrontal cortex, and the bilateral inferior parietal regions, the left superior parietal lobule, and bilateral supramarginal gyri. In addition, we found increased activity for the valid than neutral retro-cues trials in the supplementary motor area (SMA) extending into the bilateral superior and medial frontal gyri, and cingulate regions, the bilateral insula as well as the putamen. Similar patterns of the effect were also observed in the bilateral precuneus/superior parietal lobule (BA 7). Furthermore, the greater activations in the valid vs. neutral retro-cues was accompanied by lower activations in the brain regions, typically identified as default-mode network. The whole-brain analysis revealed significantly less activations for the valid than neutral retro-cue conditions in the medial prefrontal gyrus (BA 10), the anterior cingulate (BA 32) and posterior cingulate cortex (BA 31), as well as bilateral angular gyrus regions (Fig. 2A). In addition to the whole-brain analysis, we also conducted a ROI-based analysis to examine the effect of valid vs. neutral retro-cues on the independently defined brain networks established in previous literature (Dosenbach et al., 2007; Sadaghiani and D’Esposito, 2014). The results from the ROI-based analysis were consistent with the result patterns revealed in the whole-brain analysis. While both retro-cue conditions recruited the task-relevant attentional networks (i.e., fronto-parietal and cingulo-opercular ROIs; Valid: all *t* > 5.320, all *p* < 0.0001; Neutral: all *t* > 2.771, all *p* < 0.011; Fig. 2C), the corresponding linear mixed-effects model revealed that both of these networks exhibited a significantly higher activation in the valid than neutral retro-cues during the follow-up memory retention period (fronto-parietal ROI: *χ*^2^(1) = 25.171, *p* < 0.0001, ηp^2^ = 0.558; cingulo-opercular ROI: *χ*^2^(1) = 36.797, *p* < 0.0001, ηp^2^ = 0.634). As expected, the BOLD response in the default mode network was lower than baseline during memory retention for both conditions (one-sampled *t*-tests against 0: Valid: *t*(21) = –4.473, *p* = 0.00021; Neutral: *t*(21) = –2.414, *p* = 0.025), but valid retro-cues led to a significantly lower activations of this network compared to the neutral cue condition (*χ*^2^(1) = 23.481, *p* < 0.0001, ηp^2^ = 0.527).

**Table 1.** MNI coordinates and corresponding Z-scores of local maxima brain regions exhibiting a significant contrast of Valid vs. Neutral retro-cue trials during the cue and following memory retention period

| Brain region | MNI peak coordinate |  |  | Z-score | Voxels |
| --- | --- | --- | --- | --- | --- |
|  | x | y | z |  |  |
| Valid > Neutral |  |  |  |  |  |
| L Pre-SMA | −6 | 18 | 54 | 7.933 | 2639 |
| L Inferior frontal gyrus (BA 44) | −48 | 8 | 26 | 7.720 |  |
| L Dorsolateral frontal gyrus (BA 46) | −40 | 32 | 26 | 7.559 |  |
| R Dorsolateral frontal gyrus (BA 46) | 38 | 42 | 32 | 5.500 |  |
| R Middle FG (BA 6) | 44 | 2 | 54 | 5.443 |  |
| L Insula/Inferior frontal gyrus (BA 24) | −34 | 24 | 6 | 6.787 |  |
| R Insula /Inferior frontal gyrus | 36 | 24 | 2 | 7.168 | 689 |
| L Putamen | −18 | 6 | 8 | 5.758 |  |
| R Putamen | 20 | 8 | 6 | 5.162 | 54 |
| L Inferior parietal lobule (BA 40) | −36 | −52 | 50 | 7.398 | 808 |
| L Superior parietal lobule | −32 | −60 | 54 | 5.902 |  |
| L Supramarginal gyrus | −54 | −44 | 34 | 5.031 |  |
| R Inferior parietal lobule | 36 | −46 | 48 | 5.161 | 254 |
| R Supramarginal gyrus | 54 | −42 | 30 | 4.668 |  |
| R Precuneus / Superior parietal lobule | 8 | −66 | 56 | 5.935 | 96 |
| L Cerebellum | −34 | −66 | −24 | 5.016 | 103 |
| R Cerebellum | 38 | −60 | −28 | 5.886 | 202 |
| R Cerebellum (semi lunar lobule) | 24 | −70 | −48 | 4.360 | 34 |
| R Middle temporal gyrus | 54 | −34 | −4 | 4.616 | 42 |
| Neutral > Valid |  |  |  |  |  |
| R Medial frontal gyrus (BA 10) | 12 | 66 | 8 | −6.270 | 1118 |
| L Anterior cingulate (BA 32) | −4 | 26 | −10 | −6.262 |  |
| L Posterior cingulate gyrus (BA 31) / Precuneus | −6 | −46 | 32 | −5.543 | 272 |
| L Intraparietal lobule / Angular gyrus | −42 | −72 | 38 | −4.507 | 49 |
| R Angular gyrus | 54 | −66 | 36 | −4.174 | 14 |

Note: L, left; R, right; BA, Brodmann area

While the broad regions along the auditory cortex (i.e., the bilateral temporal cortex) were significantly recruited when listeners were encoding the auditory speech syllables on trial onset (Fig. S1), these regions were not recruited during memory retention. While the bilateral auditory cortical ROIs were maximally active when listeners encode the auditory syllables, activations in these regions were significant lower than baseline when listeners were maintaining syllable information in memory (Fig. 2B–C; one-sample *t*-test against 0; Valid: M = –0.094, *t*(21) = –2.247, *p* = 0.036, Neutral: M = –0.099, *t*(21) = –2.961, *p* = 0.0074).

Furthermore, these auditory cortical regions did not exhibit significantly different BOLD activation in the valid vs. neutral retro-cue trials during memory retention (Fig. 2B–C; *χ*^2^(1) = 0.088, *p* = 0.767). Likewise, the whole-brain analysis did not reveal any significant activation differences in the auditory cortex based on the cue conditions, except in the right middle temporal gyrus (BA 21) showing a relatively greater activation in the valid than neutral conditions; nevertheless, this region was not strongly active during the memory retention phase (one-sample *t*-test against 0; Valid: M = –0.0490, *t*(21) = 0.778 *p* = 0.445, Neutral: M = –0.0525, *t*(21) = 0.913, *p* = 0.372).

We further examined whether the retrocue-related modulations in the BOLD activation in the brain networks—that is, the fronto-parietal, cingulo-opercular, default-mode, and the auditory ROIs—were related to the cue-related enhancement of the perceptual precision measure (i.e., Δ ln slope *k*). The step-wise model comparison approach did not reveal any significant effect of BOLD modulations on the cue-related perceptual precision modulations; the simplest and the best model included a marginal effect of the cue-related BOLD modulation of the default-mode network ROI on the recall precision enhancement (*β* = 0.392, *t* = 1.906, *p* = 0.071) explaining 11.135% of the variance (i.e., adjusted R^2^). The leave-one-participant-out cross-validation of this model exhibited that the average model predicted outcomes did not exhibit significant relationship to the observed cue-related precision modulation (Pearson’s correlation *r* = 0.180, *p* = 0.422; average adjusted R^2^s from 22 CVs = 0.110).

### Valid retro-cues enhance neural representations of syllable category in memory

The multivariate classification analysis revealed the brain regions, in which the cued syllable category objects can be decoded from neural activities during the cue–retention phase (Fig. 3; Table 2; Table S1). Specifically, we found that neural activity patterns in the left inferior frontal gyrus/dorsolateral prefrontal cortex, an anterior portion of the left superior temporal sulcus (STS), and right precentral gyrus adjoining the right superior temporal gyrus (auditory parabelt) exhibited decoding accuracy significantly above chance when a valid retro-cue was presented. In addition, activity patterns in the right precentral region and the left parietal cortical region can accurately distinguish syllable category representations during the cue– memory retention phase. Furthermore, because retro-cues were presented visually on the screen prior to the memory retention period, we found that brain regions along the visual processing pathway, such as the left inferior temporal/fusiform gyrus region exhibited above-chance decoding accuracy for syllable category information with valid retro-cues, as expected.

**Table 2.** MNI coordinates of the task-relevant brain regions exhibiting significantly above chance decoding accuracy for auditory syllable categories

| Brain region | MNI peak coordinate |  |  |
| --- | --- | --- | --- |
|  | x | y | z |
| L Inferior frontal gyrus (BA 9) | −54 | 18 | 26 |
| R Dorsolateral prefrontal cortex | 48 | 14 | 26 |
| R Anterior inferior frontal gyrus | 50 | 42 | 8 |
| R Superior frontal gyrus | 32 | −10 | 60 |
| L Superior parietal lobule | −34 | −60 | 62 |
| L Insula | −40 | −10 | −6 |
| L Superior temporal sulcus (BA 21) | −54 | −6 | −16 |
| R Superior temporal gyrus / parabelt | 60 | 0 | 0 |
| R Postcentral gyrus | 66 | −22 | 44 |
| R Precentral gyrus / ventral premotor | 62 | 0 | 8 |
| R Angular gyrus | 60 | −52 | 36 |
| R Supramarginal gyrus | 48 | −42 | 30 |
| L Inferior temporal gyrus (BA 20) | −52 | −36 | −22 |

Note: L, left; R, right; BA, Brodmann area

However, the same analysis did not reveal any significant brain regions exhibiting above-chance decoding accuracy for syllable objects under neutral retro-cue trials. Furthermore, as expected, among the brain regions that reliably retained syllable category information (Fig. 3), none of the areas did exhibit better decoding accuracies in the neutral than valid retro-cue conditions (Δ decoding accuracies Valid – Neutral = [2.468 – 6.546%]). Note that when we performed a parametric test of the multivariate analysis against the chance level decoding (i.e., 50% against 1-tailed test), we found a qualitatively similar finding as the results derived from the non-parametric permutation-based approach (Fig. 3).

Based on the set of task-relevant neural regions that exhibited significantly accurate decoding of syllable categories during memory retention (Fig. 3; Table 2), we further examined the relationship to the behavior. Specifically, we examined whether cue-related modulation of decoding accuracy in these brain regions was related to the representational precision benefit from valid retro-cues (i.e., Δ ln slope *k*) using a stepwise multiple linear regression analysis approach (see Materials and Methods). This analysis revealed significant contributions of nine brain regions in predicting the degree of perceptual precision benefit from valid retro-cues (adjusted R^2^ = 0.492; *F*(10,11) = 3.033; *p* = 0.0413; Fig. 4A). The significant contributions of the brain regions are listed in Table 3. We found that the neural decoding accuracy modulations in the left STS, right postcentral gyrus, and right SMG exhibited significant relationships to the precision gain from valid retro-cues. We also found significant relationships to the precision benefit with the neural decoding modulations in the right dorsolateral prefrontal cortex, right precental gyrus, as well as left insula. However, we did not find any significant contributions of the inferior frontal gyri, right superior temporal gyrus, and the left inferior temporal gyrus.

**Fig. 4.**
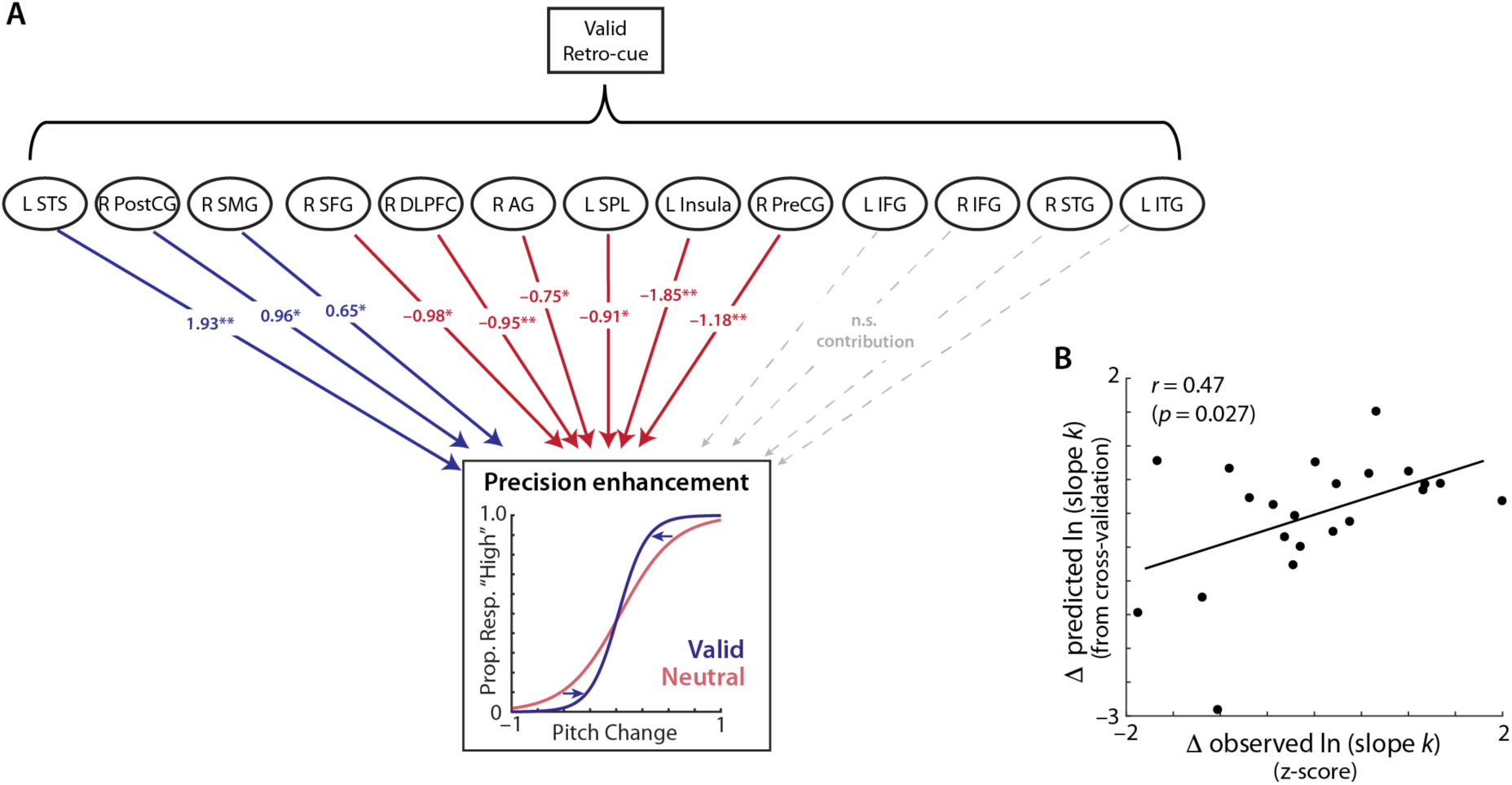
The relationship between the neural decoding for syllable category representations in memory and behavioral modulations. (**A**) Estimated contributions of the brain regions’ neural decoding modulations are resulted from the step-wise multiple linear regression model, which significantly predicted the extent of perceptual precision modulations (quantified as log- transformed slope *k* from psychophysical modeling) from Valid vs. Neutral cue conditions; * *p* < 0.05, ** *p* < 0.01, *** *p* < 0.005. (**B**) Correlation between observed and the cross-validated model- predicted cue-related modulations in perceptual precision (ln slope *k*) of the two cue conditions (Δ: Valid – Neutral). Individual’s model-predicted perceptual precision modulation obtained from cross- validation (model fitted for nth participant from remaining *n*=21 participants). All data points were z-scored prior to the model fitting.

**Table 3.** Multiple linear regression modeling result on predicting cue-related modulation of perceptual precision from the neural decoding accuracy modulations from retro-cues

| Predictors | $\beta$ | $T$ | $p$ |
| --- | --- | --- | --- |
| L Superior temporal sulcus (BA 21) | 1.926 | 3.674 | 0.0037 |
| R Postcentral gyrus | 0.965 | 3.065 | 0.011 |
| R Supramarginal gyrus | 0.647 | 2.911 | 0.014 |
| R Dorsolateral prefrontal cortex | -0.949 | -3.851 | 0.0027 |
| L Insula | -1.846 | -3.832 | 0.0028 |
| R Precentral gyrus | -1.182 | -3.486 | 0.0051 |
| R Superior frontal gyrus | -0.978 | -3.032 | 0.011 |
| L Superior parietal lobule | -0.913 | -3.029 | 0.011 |
| R Angular gyrus | -0.748 | -2.362 | 0.038 |
| R Anterior inferior frontal gyrus | -0.389 | -1.828 | 0.095 |

Note: L, left; R, right; BA, Brodmann area

Next, we confirmed the robustness of the final model using a leave-one-participant-out cross-validation to predict the left-out participant’s precision modulation (ln *k* Valid vs. ln *k* Neutral). We found that the model-predicted precision modulation had a significant relationship to the observed outcome of the cue-related precision modulation (Fig. 4B; Pearson’s correlation *r* = 0.471, *p* = 0.0270; average adjusted R^2^ (M ± SD) = 0.490 ± 0.073).

Finally, based on the model estimates, we used a Wald statistic (ZWald) to directly compare whether the neural decoding modulations of these brain regions differentially contribute to the recall precision gain from valid retro-cues (Fig. 4A; Table 3). For a given pair of model estimates, we computed a Zwald of their difference to test whether the two estimates are significantly different from each other. We found that the behavior-predicting contribution of the neural decoding modulation of the left STS was significantly different from those of all other brain regions (all ZWald > 2.127; all *p* < 0.029; FDR-adjusted), including the right dorsolateral prefrontal cortex (ZWald = 4.042; *p* = 0.00017), the left insula (ZWald = 3.883; *p* = 0.00023). We also found that the contributions of the right dorsolateral prefrontal cortex and the left insula were significantly different from each other (ZWald = 2.475; *p* = 0.012). Table S2 reports ZWald of the differences between the model estimates.

## Discussion

How can selective attention directed to an auditory object held in memory enhance the representational precision of the object? How are multiple brain networks engaged in directing attention to auditory memory objects, and how do the memory representations held in these networks contribute to enhancing the representational precision from attention? The present study investigated the role of various neocortical brain regions, including the sensory-specific and modality-general brain areas, in supporting the benefit from attention directed to working memory objects. This study utilized psychophysical modeling and a combination of univariate and multivariate analyses of fMRI data collected during an auditory retro-cueing working memory task involving speech syllables.

### Attention directed to auditory memory objects enhances recall speed and representational precision

We found that retrospective attention facilitated auditory working memory performance. Consistent with previous work demonstrating retrospective attentional benefits on memory recall (e.g., Griffin and Nobre, 2003; Lepsien et al., 2005; Lepsien and Nobre, 2007; Backer et al., 2015, 2020; Kumar et al., 2013), valid retro-cues led to significantly faster responses than uninformative neutral cues. Such enhancement of recall speed suggests that retrospective attention facilitates selective access to mnemonic representations, and prioritizes them for upcoming task demands (Backer et al., 2020; Myers et al., 2017). Furthermore, replicating the prior findings with auditory working memory (Kumar et al., 2013; Lim et al., 2015, 2018), the psychophysical modeling results demonstrate that directing attention to a specific auditory object in memory enhances the representational precision of the attended object even in the presence of background noise. Overall, our findings align with the notion that retrospective attention enhances memory representations (Lepsien et al., 2011; Rerko and Oberauer, 2013).

Even though valid retro-cues provided information orthogonal to behavioral response planning, retrospective attention facilitated memory performances. Valid cues in the current study provided information only about the upcoming probe syllable category (i.e., object-based linguistic unit), whereas the task required listeners to make decisions based on the acoustic feature (i.e., pitch) of the cued object; nevertheless, we found that valid retro-cues led to faster and more precise responses. Our findings support the object-based account of attention (Duncan, 1984; Desimone and Duncan, 1995; Alain and Arnott, 2000; Shinn-Cunningham, 2008). Sound features are integrated into a perceptual object, and attention directed to one feature also enhances the processing of other features comprising that object. Similarly, the performance benefits from valid retro-cues suggest that selective attention directed to a specific memory object enhances its representation across varying levels of information, including the lower-level features comprising that object. As we will argue below, our fMRI results indicate that retrospective auditory attention actively engages neural resources to maintain the cued objects in memory, and the distributed neural representations of the attended objects support the attentional gain in memory representational fidelity.

### Selective attention directed to memory objects engages domain-general cognitive control networks

Our univariate results revealed that retrospective attention to auditory objects held in memory recruited domain-general task control brain regions. Valid retro-cues led to significantly higher activations in broad cortical areas than neutral cue trials, including bilateral frontal and parietal cortices, as well as SMA/ACC and bilateral insula. These brain regions largely overlap with the well-known top-down networks: (*i*) the frontoparietal network—comprised of prefrontal and parietal cortices—has been implicated in executive control, including selective attention and working memory maintenance across various modalities (Rowe et al., 2000; Owen et al., 2005; Bledowski et al., 2009); (*ii*) the cingulo-opercular network, consisting of SMA/ACC and bilateral anterior insula, is known to play a role in sustained maintenance of task control and goals (Dosenbach et al., 2007; Seeley et al., 2007). Our findings align with Backer et al.’s (2020) study that re-directed listeners’ attention to semantic representations held in auditory working memory.

Prior neuroimaging work has observed increased activations in these domain-general networks with greater cognitive demands. For instance, increased activation in the cingulo-opercular network is observed when detecting unpredictable vs. predictable events (Sadaghiani and D’Esposito, 2014) or during effortful speech comprehension (Erb et al., 2013; Vaden et al., 2013). Similarly, the fronto-parietal network activity also scales with task difficulty. For example, the fronto-parietal network activity increases when more items are held in working memory and when there are increased attentional demands (Todd and Marios, 2004; Buchsbaum et al., 2005; Xu and Chun, 2006; Emrich et al., 2013; Magen et al., 2009). Based on this evidence, our results may seem counterintuitive—that is, while valid retro-cues consistently led to faster and more precise memory recall than neutral cues, we observed increased activations in these networks in the valid vs. neutral conditions.

The increased activations in these cognitive control networks suggest a potential neural mechanism by which retrospective attention facilitates memory recall: selective attention re-directed to a specific memory object does not reduce working memory load by removing unattended objects; instead, cognitive resources become reallocated to select and prioritize the attended object in memory, which result in enhancing its representational fidelity. Our interpretation is in line with the view that considers working memory as a flexible resource, which can be dynamically allocated across memory representations, and the amount of allocated resource determines memory precision (Ma et al., 2014; Bays, 2015). An alternative, but not mutually exclusive possibility can be drawn from the activity-silent working memory account (Stokes, 2015; Wolff et al., 2017; Manohar et al., 2019): selective attention redirected to an item can reactivate its representation from activity-silent storage (Lewis-Peacock et al., 2012; Sprague et al., 2016) by recruiting the general-purpose prefrontal cortical areas. And this, in turn, can rapidly restore synaptic weights of the content-specific traces of working memory (Stokes, 2015; Manohar et al., 2019).

The current results extend our prior findings. Using the very similar experimental paradigm, our prior EEG study demonstrated that valid retro-cues led to increased neural alpha (10-Hz) oscillatory power, and the extent of the increase was related to the representational precision enhancement of the cued item (Lim et al., 2015). Overall, in accordance with prior work, our results suggest that retrospective attention engages additional neural processes (Nobre et al., 2004; Griffin and Nobre, 2003; Lepsien et al., 2005; Lepsien and Nobre, 2007). This resource re-allocation is manifested in the involvement of fronto- parietal and cingulo-opercular networks for selecting and prioritizing the cued item in memory such that it is readily available for comparison with the probe (Wallis et al., 2015; Myers et al., 2017). While the prior EEG study found a direct relationship between the neural modulations and the enhancement of the representational precision of the cued syllable (Lim et al., 2015), the current study found no such relationship. The reduced number of trials in the current study (i.e., ∼45% less than Lim et al., 2015) might not have been sufficient to detect subtle differences in neural activity. It is also possible that intrinsic connectivity network ROIs might not be the critical brain regions related to auditory working memory maintenance. Future studies would be necessary to clarify how the active engagement of the neural resources allocated to the attended item in memory is related to enhancing its representational precision. Although our findings do not support the notion that retrospective attention removes unattended memory items, the mechanism by which retrospective attention benefits memory performances can depend on other factors, such as working memory load (Astle et al., 2012), cognitive resource capacity (Machizawa et al., 2012; Lim et al., 2018), or the demand exerted by attentional reorientation itself (Magen, 2009; Backer et al., 2020). We imposed a relatively low memory load (i.e., two objects), which could have led listeners to flexibly recruit the domain-general networks to enhance the representation of the cued item without removing unattended items from memory. However, it is possible that under higher working memory load, or when attentional reorientation requires cognitive resources beyond one’s cognitive capacity, retrospective attention may remove uncued items from memory (Astle et al., 2012; Machizawa et al., 2012). Also, the neural activity modulations in the fronto-parietal network depend on the amount of attentional demand directed to working memory (Magen, 2009), which could differ based on the type of information retro-cues provide (Backer et al., 2020). Future studies with directed manipulations of working memory load and attentional demands would be necessary to understand how the distinct mechanisms and underlying neural processes become instantiated to yield memory performance benefit from retrospective attention.

While the current study focused on the bilateral activations of the fronto-parietal and cingulo-opercular networks, we observed a more robust engagement of the left-lateralized neural network when listeners redirected attention to one speech object held in memory (Fig. 2A). These brain regions—the IFG, insula, premotor cortex, as well as parietal areas extending into supramarginal gyrus (SMG)—on the left hemisphere have been implicated as the articulatory network involved in speech processing (Hickok and Poeppel, 2004) and the phonological loop of working memory (e.g., Awh et al., 1996; Jonides et al., 1998). Because we did not specifically manipulate the demand for verbal rehearsal (cf. Koelsch et al., 2009), the specific role of articulatory network on attentionally enhancing mnemonic fidelity is beyond the scope of the current work; however, our results may suggest that retrospectively attended speech memory item is represented as auditory-independent articulatory or phonological information in the frontal and parietal cortical regions (Buchsbaum et al., 2005).

### Distributed neural representations of auditory memory object differentially contribute to representational precision benefit from attention

The multivariate analysis revealed that the attended syllable held in memory was represented in distributed areas of the brain. When listeners directed their attention to a particular speech syllable guided by a valid retro-cue, we found that auditory representations of the cued syllable category could be decoded from the activity patterns in both modality-general and sensory-specific brain areas (Fig. 3). These areas include lateral prefrontal and parietal cortical areas and the insula region, parts of the domain-general attention and cognitive control networks known to support both visual and auditory working memory (Duncan and Owen, 2000; Postle et al., 2000; Todd and Marios, 2004; Noyce et al., 2017; Uluç et al., 2018; Backer et al., 2020), as well as the SMG, inferior frontal gyrus, and the sensorimotor cortices. The latter are classically implicated in verbal working memory (Awh et al., 1996; Smith et al., 1998; Baddeley, 2003; Buchsbaum et al., 2005) and phonological processing (Price et al., 1997; Hartwigsen et al., 2010; Qi et al., 2019).

Beyond these brain regions commonly implicated in working memory, we also found that speech-sensitive cortical area along the left superior temporal sulcus (STS) maintained the cued syllable information during memory retention. Relatedly, the distributed neural representations of the attended syllable memory object that we observed (Fig. 3; Table 2) closely overlap with the brain regions shown to neurally represent distinct phonemes during speech perception. Prior fMRI studies have found that the left STG/STS represents distinct phonemic categories (e.g., Arsenault and Buchsbaum, 2015; Feng et al., 2018; Yi et al., 2019). Also, the inferior frontal, parietal, sensorimotor, and somatosensory regions are shown to represent phonological features when listeners encode speech sounds (e.g., Lee et al., 2012; Correia et al., 2015; Feng et al., 2021; Liebenthal et al., 2013). Thus, our results indicate that even in the absence of physical signals, attending to internal representations of speech sounds elicits very similar neural representations related to perceptual processing of speech.

These memory representations are retained across multiple brain regions as posited by the distributed account of working memory representations (Ester et al., 2015; Christophel et al., 2017).

Interestingly, our results further suggest that widely distributed memory representations do not uniformly contribute to retrospective attentional benefit in enhancing the representational precision of the cued item. The regression analysis in predicting individual differences in the memory precision gain from valid retro-cues (Δ slope *k*) revealed that the extent of individuals’ neural fidelity enhancements in the various brain regions—including the left STS, right SMG, the lateral frontal cortex, and left insula—exhibited significant relationships to the behavioral enhancement of representational precision in recalling the cued syllable in memory. However, no such relationships were apparent in the bilateral IFG or the left inferior temporal cortex (Fig. 4). This pattern indicates that the distributed memory representations across broad cortical regions work together to enhance representational precision of the attended object held in working memory; however, they do so by exerting differential contributions.

One potential explanation for our findings is that, while the multiple brain regions redundantly represent the objects held in working memory, these regions may not maintain the same level of representations of the objects. This explanation might be consistent with a view that there is a division of labor between sensory-specific and domain-general cortices maintaining working memory contents (Spitzer et al., 2014; Sreenivasan et al., 2014; Christophel et al., 2017). Accordingly, our results may suggest that the speech-sensitive area in the left STS and the language/verbal working memory related brain regions, such as the right SMG, retain both the object-level syllable category and the acoustic featural information bound to the object; in contrast, the frontal cortical regions as well as the areas along the auditory dorsal stream (e.g., IFG, insula, premotor and parietal cortex) hold auditory-independent (hence, modality-general) information about speech representations.

This possibility is in line with the functional specializations of the aforementioned brain regions. Many prior studies have demonstrated that lateral frontal areas, encompassing the DLPFC and IFG, are sensitive to between-category, but not within-category differences (e.g., Freedman et al., 2001; Myers et al., 2009). More recently, research shows similar frontal areas maintain memory representations in an abstract format, which can be generalized across sensory modalities (Spitzer et al., 2014; Uluç et al., 2018) and be flexibly transformed based on the task-relevant goals (Lee et al., 2013; Stokes et al., 2013; Long and Kuhl, 2018). Brain regions along the dorsal auditory pathway, which includes inferior frontal, insula, parietal, sensorimotor, and somatosensory regions, are also known to be part of the speech-motor network that retain the articulatory codes rather than auditory-specific information of speech (Buchsbaum et al., 2005; Hickok and Poeppel, 2007; Rauschecker and Scott, 2009). Accordingly, neural representational increase in these regions may not enhance acoustic-related representational precision as found in our model (Fig. 4).

On the contrary, the left STS has been prominently implicated in speech sound processing, categorization, and learning (Formisano et al., 2008; Leech et al., 2009; Obleser and Eisner, 2009; Mesgarani et al., 2014; Lim et al., 2019). In particular, the anterior STS/STG is known to be sensitive to tonal speech categories and speech vowels that contain talkers’ voice and/or pitch (Bachorowski and Owren, 1999; Obleser et al., 2006; Feng et al., 2018). Therefore, the left STS may be functionally suitable for retaining acoustic details of speech sounds held in memory. Accessing the auditory syllable mnemonic representations in the left STS might allow the readout of the featural-level information (e.g., pitch) of the object, which can be enhanced by attention directed to the memory object. Consistent with emerging evidence, the contribution of the speech-sensitive brain areas suggests that speech working memory is not entirely independent of sensory/auditory processing (Buchsbaum et al., 2005; Jacquemot and Scott, 2006; Perrachione et al., 2017; Scott and Perrachione, 2019). Overall, our results suggest that while both modality-general and modality-specific brain regions maintain the working memory contents (Ester et al., 2015; Bettencourt and Xu, 2016), the representations held in the distinct regions might be differentially sensitive to top-down task-related goals vs. bottom-up stimulus-specific features (Long and Kuhl, 2018), thereby exerting differential influences in the attentional enhancement of representational precision.

Note that among the brain regions retaining syllable memory objects (Fig. 3; Table 2), none of the regions did exhibit higher decoding accuracy in the neutral vs. valid retro-cue condition. We considered this pattern as evidence of neural fidelity enhancement from selective (valid) vs. non-selective (neutral) retrospective attention. However, one might question whether lower decoding accuracy in the neutral vs. valid cue condition might result from retaining a mixture of robust representations of both syllable sounds in memory rather than reflecting attentional enhancement of the cued item. If this were true, we would likely have observed higher neural activations in the above brain regions in the neutral compared to the valid condition to actively maintain strong memory representations. We tested this possibility using the extent of BOLD responses as a proxy for active representations of memory objects; the neutral condition did not lead to greater activation compared to the valid condition, such as in the left STS (e.g., mean BOLD difference in Valid – Neutral = 0.048). Thus, we conclude that this alternative possibility—maintenance of more robust representations of two syllables in the neutral compared to valid condition—is unlikely (Emrich et al., 2013).

### Potential limitations

At first glance, potential circularity in finding brain regions that retained auditory syllable memory representations seem possible. We defined these regions based on the neural activity patterns in brain areas retaining the cued object information provided by the visual valid cues. These regions might be biased to exhibit higher decoding accuracies for memory content in the valid than the neutral cue conditions. Nevertheless, the decoding accuracy in the valid cue condition cannot solely drive the predictive relationship between the extent of individuals’ neural decoding accuracy modulations (i.e., valid vs. neutral) and of behavioral (precision slope *k*) modulations. Furthermore, our main focus was on examining the relative contributions of distributed brain regions’ enhanced representational precision, which should show little effect from the overall bias that favored the valid cue condition.

The chosen retro-cues might also cause concern. Because valid retro-cues were written syllables, all activation and representation differences observed between valid and neutral cues might at least in part be byproducts of reading or imagery evoked by written text. To minimize this concern, our classification approach specifically aimed to find the brain regions maintaining the auditory traces of speech syllable objects held in memory. The classifier was trained to learn syllable sounds when they were clearly presented as auditory probes, and tested to decode the neural activity patterns during memory maintenance (i.e., no auditory input). Furthermore, the classifier was trained only on the auditory probes of the neutral cue trials to further avoid potential bias or expectations of a particular syllable informed by valid retro-cues. Of course, we cannot completely rule out the possibility that reading written syllables on valid cues itself yields reliable neural decoding. For instance, we found that the left inferior temporal gyrus (ITG), known to be sensitive to written text (Cohen et al., 2002), exhibited above-chance accuracy in decoding syllable category maintained in memory. Nevertheless, listeners must maintain the auditory-form of the speech syllables held in memory to succeed in the task. This suggests that relying on the visually presented syllable category information alone or the potential evoked imagery from the written text cannot fully explain consistent facilitation in the task performance from valid retro-cues. Supporting this possibility, the attentional enhancement of the neural representations in the left ITG, sensitive to reading texts, was not related to the extent of the cue-related enhancement of representational precision (Fig. 4).

As our main research focus was on examining the benefits of the object-based auditory retrospective attention in highlighting the attended memory representations, and because our current design was not optimal to neurally decode the exact pitch of the syllables, we performed the MVPA analysis to decode the abstracted linguistic-level of the attended speech sounds (i.e., syllable category information) instead of the low-level pitch (F0) of the syllables. While our findings suggest that high-level object-based neural representation of syllables can significantly contribute to the behavioral facilitation in recall precision of low-level pitch information (Fig. 4), they cannot fully reveal which features (i.e., higher-level vs. lower-level) underlie enhancement in neural decoding accuracy during memory maintenance. Given the fact that complex auditory objects (such as speech sounds) consist of varying levels of information, future studies are necessary to investigate whether and how different levels of auditory memory representation—low-level acoustic features vs. high-level linguistic information—are maintained in auditory working memory, and can be enhanced by selective attention.

Lastly, while we observed significant effects of retro-cues on response times and the perceptual precision measure, we did not find such effect on overall accuracy. Because we previously found retro-cues had a modest or no effect on overall accuracy (Lim et al., 2015, 2018), the lack of effect is not surprising. This might be due to the task demand for detecting very subtle changes in the pitch of the syllables, or potential variability in participants’ performances due to inherent differences in their thresholds for detecting pitch change. Future studies could employ an adaptive tracking procedure to investigate the effect of retro-cues with explicit manipulations of attentional demand that can parametrically adjust low-level acoustic changes based on each participant’s detection threshold.

## Conclusions

As demonstrated here, working memory representations across the brain, ranging from sensory-specific and domain-general cognitive-control regions differentially contribute to the retrospective attentional gain in the fidelity of memory representations. Here, we show that while re-directing attention to the relevant auditory memory objects mainly recruits domain-general cognitive-control networks, the attentional enhancement in the neural fidelity in the superior temporal sulcus and frontal cortical regions was related to the individual gain in recall precision of auditory objects from memory, with the superior temporal sulcus best predicting the attentional gain. Together with previous research on working memory, our findings provide evidence that discrete, functionally specialized brain regions collectively contribute to maintaining and attentionally enhancing working memory representations.

## Acknowledgements

This work was supported by the Max Planck Society (Max Planck research grant to JO) and the European Research Council (ERC Consolidator grant AUDADAPT, no. 646696, to JO). We thank Steven Kalinke, Sören Nikolaus, and Dunja Kunke for their help in data collection of the study. We would also like to thank Mandy Jochemko, Anke Kummer, and Nicole Pampus for their assistance at the MRI scanner.

## Author contributions

Conceptualization and Methodology: SJL, JO; Investigation: SJL, BS, LD, JL; Formal analysis: SJL, JO, CT; Visualization: SJL, JO; Writing (original draft): SJL, JO; Writing—review & editing: SJL, JO, CT, BS, LD, JL.

## Data and materials availability

All data needed to evaluate the conclusions in the paper are present in the paper and the Supplementary Materials. The raw dataset is not publicly available due to inherently identifiable nature of the data, but other data (stimuli, behavioral response data, and resulting group contrasts) are available in the OSF repository: https://osf.io/qgdn9/

## Footnotes

Note that the slope *k* measure is analogous to the inverse of response variability transformed based on a cumulative distribution function (e.g., Bays and Husain, 2008; Kumar et al., 2013; Ma et al., 2014).

## References

Alain, C., & Arnott, S. R. (2000). Selectively attending to auditory objects. Frontiers in Biosciences, 5, d202–212.

Alavash, M., Lim, S.-J., Thiel, C., Sehm, B., Deserno, L., & Obleser, J. (2018). Dopaminergic modulation of hemodynamic signal variability and the functional connectome during cognitive performance. NeuroImage, 172, 341–356. http://doi.org/10.1016/j.neuroimage.2018.01.048

Allefeld, C., Görgen, K., & Haynes, J.-D. (2016). Valid population inference for information-based imaging: From the second-level t-test to prevalence inference. NeuroImage, 141, 378–392. http://doi.org/10.1016/j.neuroimage.2016.07.040

Arsenault, J. S., Buchsbaum, B. R. (2015). Distributed neural representations of phonological features during speech perception. Journal of Neuroscience, 35(2), 634–642. https://doi.org/10.1523/JNEUROSCI.2454-14.2015

Astle, D. E., Summerfield, J., Griffin, I., & Nobre, A. C. (2012). Orienting attention to locations in mental representations. Attention, Perception & Psychophysics, 74(1), 146–162. https://doi.org/10.3758/s13414-011-0218-3

Awh, E., Jonides, J., Smith, E. E., Schumacher, E. H., Koeppe, R. A., & Katz, S. (1996). Dissociation of Storage and Rehearsal in Verbal Working Memory: Evidence From Positron Emission Tomography. Psychological Science, 7(1), 25–31. http://doi.org/10.1111/j.1467-9280.1996.tb00662.x

Bachorowski, J.-A., & Owren, M. J. (1999). Acoustic correlates of talker sex and individual talker identity are present in a short vowel segment produced in running speech. Journal of the Acoustical Society of America, 106(2), 1054–1063. http://doi.org/10.1121/1.427115

Backer, K. C., Alain, C. (2012). Orienting attention to sound object representations attenuates change deafness. Journal of Experimental Psychology: Human Perception and Performance, 38(6), 1554–1566. https://doi.org/10.1037/a0027858

Backer, K. C., Alain, C. (2014). Attention to memory: Orienting attention to sound object representations. Psychological Research, 78(3), 439–452.

Backer, K. C., Binns, M. A., & Alain, C. (2015). Neural dynamics underlying attentional orienting to auditory representations in short-term memory. Journal of Neuroscience, 35(3), 1307–1318. http://doi.org/10.1523/JNEUROSCI.1487-14.2015

Backer, K. C., Buchsbaum, B. R., & Alain, C. (2020). Orienting Attention to Short-Term Memory Representations via Sensory Modality and Semantic Category Retro-Cues. eNeuro, 7(6), ENEURO.0018-20.2020. https://doi.org/10.1523/ENEURO.0018-20.2020

Baddeley, A. (2003). Working memory: looking back and looking forward. Nature Reviews. Neuroscience, 4(10), 829–839. http://doi.org/10.1038/nrn1201

Bays, P. M., & Husain, M. (2008). Dynamic shifts of limited working memory resources in human vision. Science, 321(5890), 851–854. http://doi.org/10.1126/science.1160575

Bays P. M. (2015). Spikes not slots: noise in neural populations limits working memory. Trends in Cognitive Sciences, 19(8), 431–438. https://doi.org/10.1016/j.tics.2015.06.004

Benjamini, Y., & Hochberg, Y. (1995). Controlling the False Discovery Rate: A Practical and Powerful Approach to Multiple Testing. Journal of the Royal Statistical Society. Series B (Methodological), 57(1), 289–300. http://doi.org/10.1111/j.2517-6161.1995.tb02031.x

Bettencourt, K. C., & Xu, Y. (2016). Decoding the content of visual short-term memory under distraction in occipital and parietal areas. Nature Reviews. Neuroscience, 19(1), 150– 157. http://doi.org/10.1038/nn.4174

Bledowski, C., Rahm, B., & Rowe, J. B. (2009). What “Works” in Working Memory? Separate Systems for Selection and Updating of Critical Information. Journal of Neuroscience, 29(43), 13735–13741. http://doi.org/10.1523/JNEUROSCI.2547-09.2009

Buchsbaum, B. R., Olsen, R. K., Koch, P., & Berman, K. F. (2005). Human dorsal and ventral auditory streams subserve rehearsal-based and echoic processes during verbal working memory. Neuron, 48(4), 687–697. https://doi.org/10.1016/j.neuron.2005.09.029

Chang, C.-C., & Lin, C.-J. (2011). LIBSVM: A Library for Support Vector Machines. ACM Transactions on Intelligent Systems and Technology, 2(3), 27:1–27:27.

Christophel, T. B., Klink, P. C., Spitzer, B., Roelfsema, P. R., & Haynes, J.-D. (2017). The Distributed Nature of Working Memory. Trends in Cognitive Sciences, 21(2), 111–124. http://doi.org/10.1016/j.tics.2016.12.007

Cohen, L., Lehericy, S., Chochon, F., Lemer, C., Rivaud, S., & Dehaene, S. (2002). Language-Specific Tuning of Visual Cortex? Functional Properties of the Visual Word Form Area. Brain, 125, 1054–1069. http://doi.org/doi.org/10.1093/brain/awf094

Correia, J. M., Jansma, B. M., & Bonte, M. (2015). Decoding Articulatory Features from fMRI Responses in Dorsal Speech Regions. Journal of Neuroscience, 35(45), 15015–15025. https://doi.org/10.1523/JNEUROSCI.0977-15.2015

Cowan, N. (2001). Metatheory of storage capacity limits. Behavioral and Brain Sciences, 24(01), 154–176.

Cox, R. W. (1996). AFNI: software for analysis and visualization of functional magnetic resonance neuroimages. Computers and Biomedical Research, an International Journal, 29(3), 162–173.

Desimone, R., & Duncan, J. (1995). Neural Mechanisms of Selective Visual Attention. Annual Review of Neuroscience, 18(1), 193–222. http://doi.org/10.1146/annurev.ne.18.030195.001205

Dosenbach, N. U. F., Fair, D. A., Cohen, A. L., Schlaggar, B. L., & Petersen, S. E. (2008). A dual-networks architecture of top-down control. Trends in Cognitive Sciences, 12(3), 99–105. http://doi.org/10.1016/j.tics.2008.01.001

Dosenbach, N. U. F., Fair, D. A., Miezin, F. M., Cohen, A. L., Wenger, K. K., Dosenbach, R. A. T., et al. (2007). Distinct brain networks for adaptive and stable task control in humans. Proceedings of the National Academy of Sciences, 104(26), 11073–11078.

Duncan, J. (1984). Selective Attention and the Organization of Visual Information. Journal of Experimental Psychology. General, 113(4), 51–517.

Duncan, J., & Owen, A. M. (2000). Common regions of the human frontal lobe recruited by diverse cognitive demands. Trends in Neurosciences, 23(10), 475–483. http://doi.org/10.1016/S0166-2236(00)01633-7

Emrich, S. M., Riggall, A. C., LaRocque, J. J., & Postle, B. R. (2013). Distributed Patterns of Activity in Sensory Cortex Reflect the Precision of Multiple Items Maintained in Visual Short-Term Memory. Journal of Neuroscience, 33(15), 6516–6523. http://doi.org/10.1523/JNEUROSCI.5732-12.2013

Erb, J., Henry, M. J., Eisner, F., & Obleser, J. (2013). The brain dynamics of rapid perceptual adaptation to adverse listening conditions. Journal of Neuroscience, 33(26), 10688–10697. http://doi.org/10.1523/JNEUROSCI.4596-12.2013

Ester, E. F., Sprague, T. C., & Serences, J. T. (2015). Parietal and Frontal Cortex Encode Stimulus-Specific Mnemonic Representations during Visual Working Memory. Neuron, 87(4), 893–905. http://doi.org/10.1016/j.neuron.2015.07.013

Fedorenko, E., Behr, M. K., & Kanwisher, N. (2011). Functional specificity for high-level linguistic processing in the human brain. Proceedings of the National Academy of Sciences, 108(39), 16428–16433. http://doi.org/10.1073/pnas.1112937108

Feng, G., Yi, H.-G., & Chandrasekaran, B. (2018). The Role of the Human Auditory Corticostriatal Network in Speech Learning. Cerebral Cortex, 29(10), 4077–4089. http://doi.org/10.1093/cercor/bhy289

Feng, G., Gan, Z., Yi, H.-G., Ell, S. W., Roark, C. L., Wang, S., Wong, P. C. M, & Chandrasekaran, B. (2021). Neural dynamics underlying the acquisition of distinct auditory category structures, NeuroImage, 244(1), 118565. https://doi.org/10.1016/j.neuroimage.2021.118565

Formisano, E., De Martino, F., Bonte, M., & Goebel, R. (2008). “Who” is saying “what?” Brain-based decoding of human voice and speech. Science, 322(5903), 970–973. http://doi.org/10.1126/science.1164318

Freedman, D. J., Riesenhuber, M., Poggio, T., & Miller, E. K. (2001). Categorical representation of visual stimuli in the primate prefrontal cortex. Science, 291(5502), 312–316. http://doi.org/10.1126/science.291.5502.312

Fritz, J. B., Elhilali, M., David, S. V., & Shamma, S. A. (2007). Auditory attention — focusing the searchlight on sound. Current Opinion in Neurobiology, 17(4), 437–455. http://doi.org/10.1016/j.conb.2007.07.011

Gazzaley, A., & Nobre, A. C. (2012). Top-down modulation: bridging selective attention and working memory. Trends in Cognitive Sciences, 16(2), 129–135. http://doi.org/10.1016/j.tics.2011.11.014

Goldman-Rakic, P. S. (1995). Cellular Basis of Working Memory Review. Neuron, 14, 477– 485.

Griffin, I. C., & Nobre, A. C. (2003). Orienting Attention to Locations in Internal Representations. Journal of Cognitive Neuroscience, 15(8), 1176–1194.

Harrison, S. A., & Tong, F. (2009). Decoding reveals the contents of visual working memory in early visual areas. Nature, 458(7238), 632–635. http://doi.org/10.1038/nature07832

Hartwigsen, G., Baumgaertner, A., Price, C. J., Koehnke, M., Ulmer, S., & Siebner, H. R. (2010). Phonological decisions require both the left and right supramarginal gyri. Proceedings of the National Academy of Sciences, 107(38), 16494–16499. http://doi.org/10.1073/pnas.1008121107

Haynes, J.-D., Sakai, K., Rees, G., Gilbert, S., Frith, C., Passingham, R. E. (2007). Reading hidden intentions in the human brain. Current Biology 17, 323–328.

Herbst, S. K., & Obleser, J. (2019). Implicit temporal predictability enhances pitch discrimination sensitivity and biases the phase of delta oscillations in auditory cortex. NeuroImage, 203, 116198. https://doi.org/10.1016/j.neuroimage.2019.116198

Hickok, G., & Poeppel, D. (2007). The cortical organization of speech processing. Nature Reviews. Neuroscience, 8(5), 393–402. http://doi.org/10.1038/nrn2113

Higo, T., Mars, R. B., Boorman, E. D., Buch, E. R., & Rushworth, M. F. S. (2011). Distributed and causal influence of frontal operculum in task control. Proceedings of the National Academy of Sciences of the United States of America, 108(10), 4230–4235. http://doi.org/10.1073/pnas.1013361108

Jacquemot, C., & Scott, S. K. (2006). What is the relationship between phonological short-term memory and speech processing? Trends in Cognitive Sciences, 10(11), 480–486. http://doi.org/10.1016/j.tics.2006.09.002

Johnson, M. K., Raye, C. L., Mitchell, K. J., Greene, E. J., Cunningham, W. A., Sanislow, C. A. (2005). Using fMRI to investigate a component process of reflection: prefrontal correlates of refreshing a just-activated representation. Cognitive, affective & behavioral neuroscience, 5(3), 339–361. https://doi.org/10.3758/cabn.5.3.339

Jonides, J., Schumacher, E. H., Smith, E. E., Koeppe, R. A., Awh, E., Reuter-Lorenz, P. A., Marshuetz, C., Willis, C. R. (1998). The Role of Parietal Cortex in Verbal Working Memory. The Journal of Neuroscience, 18(13), 5026–5034.

Koelsch, S., Schulze, K., Sammler, D., Fritz, T., Müller, K., & Gruber, O. (2009). Functional architecture of verbal and tonal working memory: an FMRI study. Human Brain Mapping, 30, 859–873. http://doi.org/10.1002/hbm.20550’

Kriegeskorte, N., Goebel, R., & Bandettini, P. (2006). Information-based functional brain mapping. Proceedings of the National Academy of Sciences, 103(10), 3863–3868. http://doi.org/10.1073/pnas.0600244103

Kumar S, Joseph S, Pearson B, Teki S, Fox ZV, Griffiths TD, Husain M. (2013). Resource allocation and prioritization in auditory working memory. Cognitive Neuroscience, 4:12– 20. doi: 10.1080/17588928.2012.716416.

Kumar, S., Joseph, S., Gander, P. E., Barascud, N., Halpern, A. R., & Griffiths, T. D. (2016). A Brain System for Auditory Working Memory. Journal of Neuroscience, 36(16), 4492– 4505. http://doi.org/10.1523/JNEUROSCI.4341-14.2016

Kuo, B.-C., Stokes, M. G., & Nobre, A. C. (2012). Attention modulates maintenance of representations in visual short-term memory. Journal of Cognitive Neuroscience, 24(1), 51–60. http://doi.org/10.1162/jocn_a_00087

Lee, S.-H., Kravitz, D. J., & Baker, C. I. (2013). Goal-dependent dissociation of visual and prefrontal cortices during working memory. Nature Neuroscience, 16(8), 997–999. http://doi.org/10.1038/nn.3452

Lee, Y. S., Turkeltaub, P., Granger, R., & Raizada, R. D. (2012). Categorical speech processing in Broca’s area: an fMRI study using multivariate pattern-based analysis. Journal of Neuroscience, 32(11), 3942–3948.

Leech, R., Holt, L. L., Devlin, J. T., & Dick, F. (2009). Expertise with artificial nonspeech sounds recruits speech-sensitive cortical regions. Journal of Neuroscience, 29(16), 5234–5239. http://doi.org/10.1523/JNEUROSCI.5758-08.2009

Lepsien, J., Nobre, A. C. (2006). Cognitive control of attention in the human brain: Insights from orienting attention to mental representations. Brain Research, 1105(1), 20–31. https://doi.org/10.1016/j.brainres.2006.03.033

Lepsien, J., & Nobre, A. C. (2007). Attentional Modulation of Object Representations in Working Memory. Cerebral Cortex, 17(9), 2072–2083. http://doi.org/10.1093/cercor/bhl116

Lepsien, J., Griffin, I. C., Devlin, J. T., & Nobre, A. C. (2005). Directing spatial attention in mental representations: Interactions between attentional orienting and working-memory load. NeuroImage, 26(3), 733–743. http://doi.org/10.1016/j.neuroimage.2005.02.026

Lepsien, J., Thornton, I., & Nobre, A. C. (2011). Modulation of working-memory maintenance by directed attention. Neuropsychologia, 49(6), 1569–1577. http://doi.org/10.1016/j.neuropsychologia.2011.03.011

Lewis-Peacock, J. A., Drysdale, A. T., Oberauer, K., & Postle, B. R. (2012). Neural Evidence for a Distinction between Short-term Memory and the Focus of Attention. Journal of Cognitive Neuroscience, 24(1), 61–79. http://doi.org/10.1162/jocn_a_00140

Liebenthal, E., Sabri, M., Beardsley, S. A., Mangalathu-Arumana, J., Desai, A. (2013). Neural Dynamics of Phonological Processing in the Dorsal Auditory Stream. Journal of Neuroscience, 33(39), 15414–15424.

Lim, S.-J., Fiez, J. A., & Holt, L. L. (2019). Role of the striatum in incidental learning of sound categories. Proceedings of the National Academy of Sciences, 1–10. http://doi.org/10.1073/pnas.1811992116

Lim, S.-J., Wöstmann, M., & Obleser, J. (2015). Selective Attention to Auditory Memory Neurally Enhances Perceptual Precision. Journal of Neuroscience, 35(49), 16094– 16104. http://doi.org/10.1523/JNEUROSCI.2674-15.2015

Lim, S.-J., Wöstmann, M., Geweke, F., & Obleser, J. (2018). The Benefit of Attention-to- Memory Depends on the Interplay of Memory Capacity and Memory Load. Frontiers in Psychology, 9, 146. http://doi.org/10.3389/fpsyg.2018.00184

Linke, A. C., & Cusack, R. (2015). Flexible Information Coding in Human Auditory Cortex during Perception, Imagery, and STM of Complex Sounds. Journal of Cognitive Neuroscience, 27(7), 1322–1333. http://doi.org/10.1162/jocn_a_00780

Linke, A. C., Vicente-Grabovetsky, A., & Cusack, R. (2011). Stimulus-specific suppression preserves information in auditory short-term memory. Proceedings of the National Academy of Sciences of the United States of America, 108(31), 12961–12966. http://doi.org/10.1073/pnas.1102118108

Long, N. M., & Kuhl, B. A. (2018). Bottom-up and top-down factors differentially influence stimulus representations across large-scale attentional networks. Journal of Neuroscience, 38(10), 2724–17–2504. http://doi.org/10.1523/JNEUROSCI.2724-17.2018

Luck, S. J., & Vogel, E. K. (2013). Visual working memory capacity: from psychophysics and neurobiology to individual differences. Trends in Cognitive Sciences, 17(8), 391–400. http://doi.org/10.1016/j.tics.2013.06.006

Ma, W. J., Husain, M., & Bays, P. M. (2014). Changing concepts of working memory. Nature Reviews. Neuroscience, 17(3), 347–356. http://doi.org/10.1038/nn.3655

Machizawa, M. G., Goh, C. C. W., Driver, J. (2012). Human Visual Short-Term Memory Precision Can Be Varied at Will When the Number of Retained Items Is Low. Psychological Science, 23(6), 554–559. https://doi.org/10.1177/0956797611431988

Magen, H., Emmanouil, T. A., McMains, S. A., Kastner, S., & Treisman, A. (2009). Attentional demands predict short-term memory load response in posterior parietal cortex. Neuropsychologia, 47(8-9), 1790–1798. https://doi.org/10.1016/j.neuropsychologia.2009.02.015

Manohar, S. G., Zokaei, N., Fallon, S. J., Vogels, T. P., & Husain, M. (2019). Neural mechanisms of attending to items in working memory. Neuroscience and Biobehavioral Reviews, 101, 1–12. http://doi.org/10.1016/j.neubiorev.2019.03.017

Mesgarani, N., Cheung, C., Johnson, K., & Chang, E. F. (2014). Phonetic feature encoding in human superior temporal gyrus. Science, 343(6174), 1006–1010. http://doi.org/10.1126/science.1245994

Misaki, M., Kim, Y., Bandettini, P. A., & Kriegeskorte, N. (2010). Comparison of multivariate classifiers and response normalizations for pattern-information fMRI. NeuroImage, 53(1), 103–118. http://doi.org/10.1016/j.neuroimage.2010.05.051

Murray, A. M., Nobre, A. C., Clark, I. A., Cravo, A. M., & Stokes, M. G. (2013). Attention Restores Discrete Items to Visual Short-Term Memory. Psychological Science, 24(4), 550–556. http://doi.org/10.1177/0956797612457782

Myers, E. B., Blumstein, S. E., Walsh, E., & Eliassen, J. (2009). Inferior frontal regions underlie the perception of phonetic category invariance. Psychological Science, 20(7), 895–903. http://doi.org/10.1111/j.1467-9280.2009.02380.x

Myers, N. E., Stokes M. G., Walther L., Nobre A. C., (2014). Oscillatory brain state predicts variability in working memory. Journal of Neuroscience, 34(23):7735–43. doi: 10.1523/JNEUROSCI.4741-13.2014.

Myers, N. E., Stokes, M. G., & Nobre, A. C. (2017). Prioritizing Information during Working Memory: Beyond Sustained Internal Attention. Trends in Cognitive Sciences, 21(6), 449–461. http://doi.org/10.1016/j.tics.2017.03.010

Nobre, A. C., Coull, J. T., Maquet, P., Frith, C. D., van den Berg, R., Mesulam, M. M. (2004). Orienting Attention to Locations in Perceptual Versus Mental Representations. Journal of Cognitive Neuroscience, 16(3), 363–373. https://doi.org/10.1162/089892904322926700

Noyce, A. L., Cestero, N., Michalka, S. W., Shinn-Cunningham, B. G., & Somers, D. C. (2017). Sensory-Biased and Multiple-Demand Processing in Human Lateral Frontal Cortex. Journal of Neuroscience, 37(36), 8755–8766. http://doi.org/10.1523/JNEUROSCI.0660-17.2017

Obleser, J., & Eisner, F. (2009). Pre-lexical abstraction of speech in the auditory cortex. Trends in Cognitive Sciences, 13(1), 14–19. http://doi.org/10.1016/j.tics.2008.09.005

Obleser, J., Boecker, H., Drzezga, A., Haslinger, B., Hennenlotter, A., Roettinger, M., et al. (2006). Vowel sound extraction in anterior superior temporal cortex. Human Brain Mapping, 27(7), 562–571. http://doi.org/10.1002/hbm.20201

Owen, A. M., McMillan, K. M., Laird, A. R., & Bullmore, E. (2005). N-back working memory paradigm: A meta-analysis of normative functional neuroimaging studies. Human Brain Mapping, 25(1), 46–59. http://doi.org/10.1002/hbm.20131

Perrachione, T. K., Ghosh, S. S., Ostrovskaya, I., Gabrieli, J. D. E., & Kovelman, I. (2017). Phonological Working Memory for Words and Nonwords in Cerebral Cortex. *Journal of Speech*, Language, and Hearing Research, 60(7), 1959–1979. http://doi.org/10.1044/2017_JSLHR-L-15-0446

Postle, B. R. (2006). Working memory as an emergent property of the mind and brain. Neuroscience, 139(1), 23–38. http://doi.org/10.1016/j.neuroscience.2005.06.005

Postle, B. R., Stern, C. E., Rosen, B. R., & Corkin, S. (2000). An fMRI Investigation of Cortical Contributions to Spatial and Nonspatial Visual Working Memory. NeuroImage, 11(5), 409–423. http://doi.org/10.1006/nimg.2000.0570

Power, J. D., Cohen, A. L., Nelson, S. M., Wig, G. S., Barnes, K. A., Church, J. A., et al. (2011). Functional Network Organization of the Human Brain. Neuron, 72(4), 665–678. http://doi.org/10.1016/j.neuron.2011.09.006

Price, C. J., Moore, C. J., Humphreys, G. W., & Wise, R. J. S. (1997). Segregating semantic from phonological processes during reading. Journal of Cognitive Neuroscience, 9(6), 727–733. http://doi.org/10.1162/jocn.1997.9.6.727

Qi, Z., Han, M., Wang, Y., de Los Angeles, C., Liu, Q., Garel, K., et al. (2019). Speech processing and plasticity in the right hemisphere predict variation in adult foreign language learning. NeuroImage, 192, 76–87. http://doi.org/10.1016/j.neuroimage.2019.03.008

Raichle, M. E., MacLeod, A. M., Snyder, A. Z., Powers, W. J., Gusnard, D. A., & Shulman, G. L. (2001). A default mode of brain function. Proceedings of the National Academy of Sciences, 98(2), 676–682.

Rauschecker, J. P., Scott, S. K. (2009). Maps and streams in the auditory cortex: Nonhuman primates illuminate human speech processing. Nature Neuroscience, 12(6), 718–724. https://doi.org/10.1038/nn.2331

Rerko, L., & Oberauer, K. (2013). Focused, unfocused, and defocused information in working memory. *Journal of Experimental Psychology: Learning*, Memory, and Cognition, 39(4), 1075–1096. http://doi.org/10.1037/a0031172

Riggall, A. C., & Postle, B. R. (2012). The Relationship between Working Memory Storage and Elevated Activity as Measured with Functional Magnetic Resonance Imaging. Journal of Neuroscience, 32(38), 12990–12998. http://doi.org/10.1523/JNEUROSCI.1892-12.2012

Rohenkohl, G., Cravo, A. M., Wyart, V., & Nobre, A. C. (2012). Temporal expectation improves the quality of sensory information. Journal of Neuroscience, 32(24), 8424– 8428. https://doi.org/10.1523/JNEUROSCI.0804-12.2012

Rowe, J. B., Toni, I., Josephs, O., Frackowiak, R. S., & Passingham, R. E. (2000). The prefrontal cortex: response selection or maintenance within working memory? Science, 288(5471), 1656–1660.

Sadaghiani, S., & D’Esposito, M. (2014). Functional Characterization of the Cingulo- Opercular Network in the Maintenance of Tonic Alertness. Cerebral Cortex, 1–11. http://doi.org/10.1093/cercor/bhu072

Scott, T. L., & Perrachione, T. K. (2019). Common cortical architectures for phonological working memory identified in individual brains. NeuroImage, 202, 116096. http://doi.org/10.1016/j.neuroimage.2019.116096

Seeley, W. W., Menon, V., Schatzberg, A. F., Keller, J., Glover, G. H., Kenna, H., et al. (2007). Dissociable Intrinsic Connectivity Networks for Salience Processing and Executive Control. Journal of Neuroscience, 27(9), 2349–2356. http://doi.org/10.1523/JNEUROSCI.5587-06.2007

Serences, J. T., Ester, E. F., Vogel, E. K., & Awh, E. (2009). Stimulus-specific delay activity in human primary visual cortex. Psychological Science, 20(2), 207–214. http://doi.org/10.1111/j.1467-9280.2009.02276.x

Shinn-Cunningham, B. G. (2008). Object-based auditory and visual attention. Trends in Cognitive Sciences, 12(5), 182–186. http://doi.org/10.1016/j.tics.2008.02.003

Smith, E. E., Jonides, J., Marshuetz, C., & Koeppe, R. A. (1998). Components of verbal working memory: Evidence fromneuroimaging. Proceedings of the National Academy of Sciences, 95(3), 876–882. http://doi.org/10.1073/pnas.95.3.876

Souza, A. S., & Oberauer, K. (2016). In search of the focus of attention in working memory: 13 years of the retro-cue effect. *Attention, Perception*, & Psychophysics, 78(7), 1839– 1860. http://doi.org/10.3758/s13414-016-1108-5

Souza, A. S., Rerko, L., & Oberauer, K. (2014). Unloading and reloading working memory: Attending to one item frees capacity. Journal of Experimental Psychology. Human Perception and Performance, 40(3), 1237–1256. http://doi.org/10.1037/a0036331

Spitzer, B., Fleck, S., & Blankenburg, F. (2014). Parametric Alpha- and Beta-Band Signatures of Supramodal Numerosity Information in Human Working Memory. Journal of Neuroscience, 34(12), 4293–4302. http://doi.org/10.1523/JNEUROSCI.4580-13.2014

Sprague, T. C., Ester, E. F., & Serences, J. T. (2016). Restoring Latent Visual Working Memory Representations in Human Cortex. Neuron, 91(3), 694–707. http://doi.org/10.1016/j.neuron.2016.07.006

Sreenivasan, K. K., Gratton, C., Vytlacil, J., & D’Esposito, M. (2014). Evidence for working memory storage operations in perceptual cortex. *Cognitive*, Affective & Behavioral Neuroscience, 14(1), 117–128. http://doi.org/10.3758/s13415-013-0246-7

Stelzer, J., Chen, Y., & Turner, R. (2013). Statistical inference and multiple testing correction in classification-based multi-voxel pattern analysis (MVPA): Random permutations and cluster size control. NeuroImage, 65, 69–82. http://doi.org/10.1016/j.neuroimage.2012.09.063

Stokes, M. G. (2015). “Activity-silent” working memory in prefrontal cortex: a dynamic coding framework. Trends in Cognitive Sciences, 19(7), 394–405. http://doi.org/10.1016/j.tics.2015.05.004

Stokes, M. G., Kusunoki, M., Sigala, N., Nili, H., Gaffan, D., & Duncan, J. (2013). Dynamic Coding for Cognitive Control in Prefrontal Cortex. Neuron, 78(2), 364–375. http://doi.org/10.1016/j.neuron.2013.01.039

Todd, J. J., & Marios, R. (2004). Capacity limit of visual short-term memory in human posterior parietal cortex. Nature Neuroscience, 428, 751–754.

Uluç, I., Schmidt, T. T., Wu, Y.-H., & Blankenburg, F. (2018). Content-specific codes of parametric auditory working memory in humans. NeuroImage, 183, 254–262.

Vaden, K. I., Kuchinsky, S. E., Cute, S. L., Ahlstrom, J. B., Dubno, J. R., & Eckert, M. A. (2013). The Cingulo-Opercular Network Provides Word-Recognition Benefit. Journal of Neuroscience, 33(48), 18979–18986. http://doi.org/10.1523/JNEUROSCI.1417-13.2013

Wallis, G., Stokes, M., Cousijn, H., Woolrich, M., & Nobre, A. C. (2015). Frontoparietal and Cingulo-opercular Networks Play Dissociable Roles in Control of Working Memory. Journal of Cognitive Neuroscience, 15, 1–16. http://doi.org/10.1162/jocn_a_00838

Wilsch, A., & Obleser, J. (2016). What works in auditory working memory? A neural oscillations perspective. Brain Research, 1640 (Part B), 193–207. http://doi.org/10.1016/j.brainres.2015.10.054

Wolff, M. J., Jochim, J., Akyürek, E. G., & Stokes, M. G. (2017). Dynamic hidden states underlying working-memory-guided behavior. Nature Reviews. Neuroscience, 20(6), 864–871. http://doi.org/10.1038/nn.4546

Xu, Y., & Chun, M. M. (2006). Dissociable neural mechanisms supporting visual short-term memory for objects. Nature, 440(7080), 91–95. http://doi.org/10.1038/nature04262

Yi, H. G., Leonard, M. K., & Chang, E. F. (2019). The Encoding of Speech Sounds in the Superior Temporal Gyrus. Neuron, 102(6), 1096–1110. https://doi.org/10.1016/j.neuron.2019.04.023

Zhang, W., & Luck, S. J. (2008). Discrete fixed-resolution representations in visual working memory. Nature, 453(7192), 233–235. http://doi.org/10.1038/nature06860

Zimmermann, J. F., Moscovitch, M., Alain, C. (2016). Attending to auditory memory. Brain Research, 1640, 208–221. https://doi.org/10.1016/j.brainres.2015.11.032

