## Supplementary Materials for "Distributed networks for auditory memory differentially contribute to recall precision"

Sung-Joo Lim*, Christiane Thiel, Bernhard Sehm, Lorenz Deserno,

Jöran Lepsien, and Jonas Obleser*

**This PDF file includes:**

Fig. S1

Tables S1 to S2

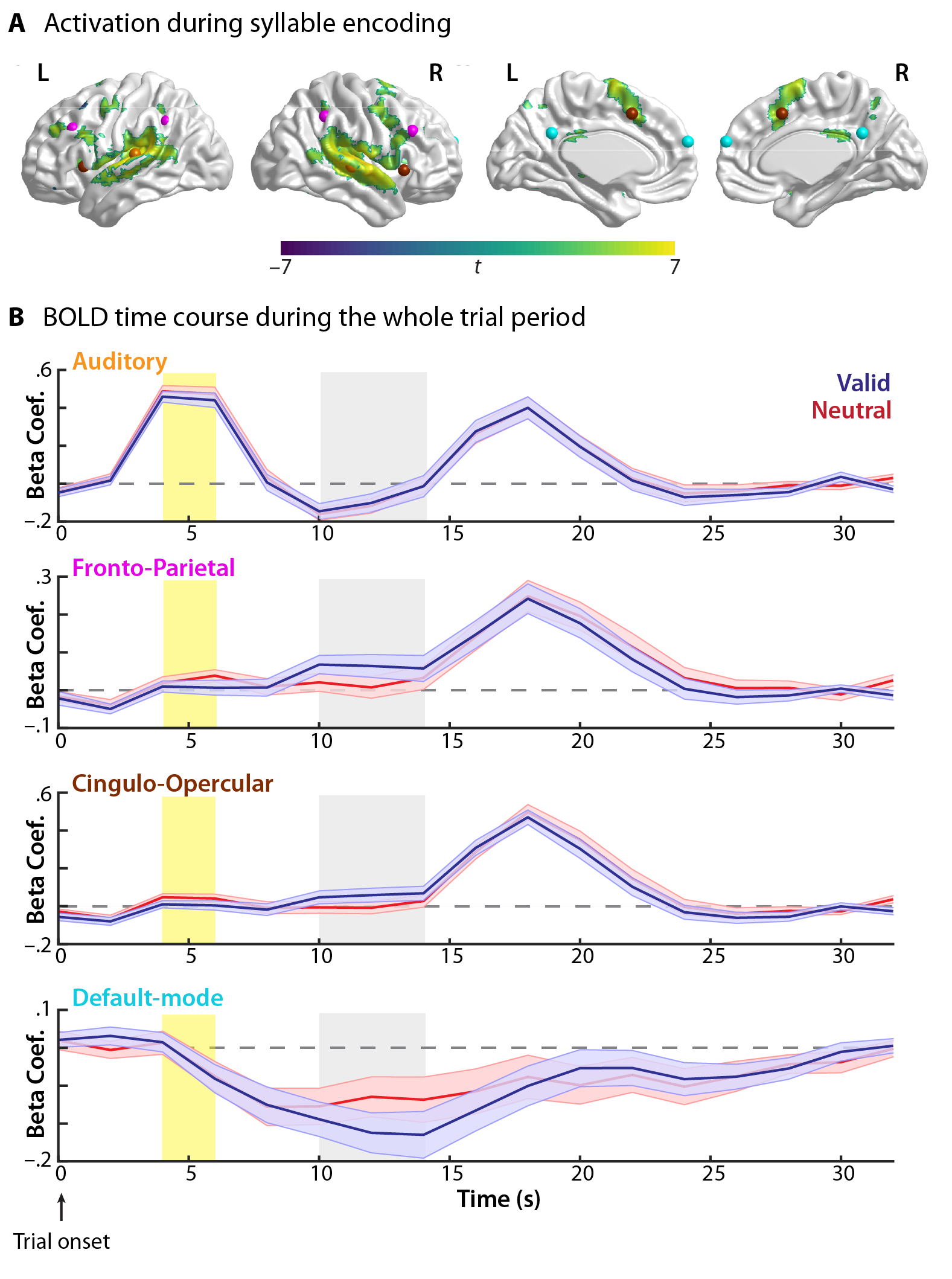
**Fig. S1.** Neural activation during the syllable encoding phase and throughout performing the auditory working memory task. (A) Brain clusters significantly active when listeners encode two syllable sounds at the onset of the trial (collapsed for both cue conditions; thresholded at *p* < 0.001 and corrected for multiple comparisons at *p* < .05). The ROIs are indicated with spheres of the corresponding colors (as shown in B and see Methods and Materials). L, left; R, right. (B) BOLD time courses during performing the entire task trial extracted from the corresponding ROIs, for illustration purpose only. We used a general linear model (GLM) with finite-impulse response (FIR) basis functions (0–32 s relative to trial onset) to illustrate BOLD activity during the entire trial of the task in the valid and neutral conditions. The yellow shaded time points represent the neural activation capturing the syllable encoding phase of the task; the grey shaded time points capture the BOLD activation related to memory retention following retro-cues. Dashed lines indicate no activation (beta-coefficient = 0).

Table S1. MNI coordinates of clusters in the whole brain exhibiting significantly above chance decoding accuracy for auditory syllable categories

| Brain region | MNI peak coordinate | | |
| --- | --- | --- | --- |
|  | *x* | *y* | *z* |
| L Inferior frontal gyrus (BA 9) | –54 | 18 | 26 |
| R Dorsolateral prefrontal cortex | 48 | 14 | 26 |
| R Anterior inferior frontal gyrus | 50 | 42 | 8 |
| R Superior frontal gyrus | 32 | –10 | 60 |
| L Superior parietal lobule | –34 | –60 | 62 |
| L Insula | –40 | –10 | –6 |
| L Superior temporal sulcus (BA 21) | –54 | –6 | –16 |
| R Heschl’s gyrus | 60 | 0 | 0 |
| R Postcentral gyrus | 66 | –22 | 44 |
| R Precental gyrus | 62 | 0 | 8 |
| R Angular gyrus | 60 | –52 | 36 |
| R Supramarginal gyrus | 48 | –42 | 30 |
| L Inferior temporal gyrus (BA 20) | –52 | –36 | –22 |
| R Superior medial frontal gyrus | 12 | 60 | 21 |
| L Medial frontal gyrus | –9 | 36 | –18 |
| L Anterior medial prefrontal cortex | –9 | 66 | 18 |
| R Middle cingulate gyrus | 18 | 21 | 36 |
| L Middle cingulate gyrus | –12 | –6 | 33 |
| L Parahippocampal | –24 | –36 | 6 |
| L Cuneus | –15 | –72 | 15 |
| R Frontal Pole | 21 | 54 | –12 |
| L Frontal Pole | –36 | 48 | –9 |
| R Cerebellum | 3 | –72 | –33 |
| R Cerebellum | 51 | –51 | –45 |
| R Cerebellum / Crus II | 18 | –75 | –42 |

Note: L, left; R, right; BA, Brodmann area

Table S2. Wald Z statistic and *p* values of the pairwise comparison of the contributions of the neural decoding modulations of the brain regions to recall precision gain from valid retro-cues

|  | R PostCG | R SMG | R DLPFC | L Insula | R PreCG | R SFG | L SPL | R AG | R IFG |
| --- | --- | --- | --- | --- | --- | --- | --- | --- | --- |
| L STS | 2.13* | 2.60* | 4.04* | 3.88* | 4.22* | 3.76* | 3.92* | 3.45* | 3.64* |
| R PostCG |  | 0.90 | 4.50* | 4.03* | 3.57* | 3.38* | 3.47* | 3.21* | 2.94* |
| R SMG |  |  | 4.25* | 4.09* | 4.51* | 3.62* | 4.00* | 2.89* | 3.03* |
| R DPLFC |  |  |  | 2.47* | 0.64 | 0.079 | 0.12 | 0.60 | 1.73 |
| L Insula |  |  |  |  | 1.71 | 1.91* | 2.30* | 2.92* | 3.02* |
| R PreCG |  |  |  |  |  | 0.49 | 1.15 | 1.02 | 2.12* |
| R SFG |  |  |  |  |  |  | 0.16 | 0.85 | 2.38* |
| L SPL |  |  |  |  |  |  |  | 0.39 | 1.48 |
| R AG |  |  |  |  |  |  |  |  | 1.20 |

Note: Upper diagonal values are listed. L: Left; R: Right; STS: superior temporal sulcus; PostCG: postcentral gyrus; SMG: supramarginal gyrus; DLPFC: dorsolateral prefrontal cortex; PreCG: precentral gyrus; SFG: superior frontal gyrus; SPL: superior parietal lobule; AG: angular gyrus; IFG: inferior frontal gyrus. **p* < 0.05 (FDR-adjusted).
